# Selection and gene flow define polygenic barriers between incipient butterfly species

**DOI:** 10.1101/2020.04.09.034470

**Authors:** Steven M. Van Belleghem, Jared M. Cole, Gabriela Montejo-Kovacevich, Caroline N. Bacquet, W. Owen McMillan, Riccardo Papa, Brian A. Counterman

## Abstract

Characterizing the genetic architecture of species boundaries remains a difficult task. Hybridizing species provide a powerful system to identify the factors that shape genomic variation and, ultimately, identify the regions of the genome that maintain species boundaries. Unfortunately, complex histories of isolation, admixture and selection can generate heterogenous genomic landscapes of divergence which make inferences about the regions that are responsible for species boundaries problematic. However, as the signal of admixture and selection on genomic loci varies with recombination rate, their relationship can be used to infer their relative importance during speciation. Here, we explore patterns of genomic divergence, admixture and recombination rate among hybridizing lineages across the *Heliconius erato* radiation. We focus on the incipient species, *H. erato* and *H. himera*, and distinguish the processes that drive genomic divergence across three contact zones where they frequently hybridize. Using demographic modeling and simulations, we infer that periods of isolation and selection have been major causes of genome-wide correlation patterns between recombination rate and divergence between these incipient species. Upon secondary contact, we found surprisingly highly asymmetrical introgression between the species pair, with a paucity of *H. erato* alleles introgressing into the *H. himera* genomes. We suggest that this signal may result from a current polygenic species boundary between the hybridizing lineages. These results contribute to a growing appreciation for the importance of polygenic architectures of species boundaries and pervasive genome-wide selection during the early stages of speciation with gene flow.

## Introduction

Disentangling the factors that drive genomic divergence is necessary for advancing our understanding of speciation. Targets of selection, for example, may be responsible for adaptive differences between species; and their number, distribution and effect on gene flow along the genome define the architecture of species boundaries. In population genomic studies, targets of selection are expected to show elevated divergence between species and reduced genetic diversity within species [1]. These highly divergent loci often reflect local adaptation and/or incompatibilities between species, and can be considered the loci that define the species [2,3]. This is because natural selection acts as a local genomic “barrier” to gene flow between hybridizing species [4–6]. In contrast, the rest of the genome, which is not under such selective pressures, may be expected to exchange more freely between the species (i.e. admixture). However, genetic variation at these latter genomic regions can be greatly impacted by neutral demographic processes (i.e. population size and migration) and the indirect effects of nearby targets of selection (i.e. linked selection). Local recombination rates can further influence how these processes impact genetic variation, which collectively result in highly heterogenous patterns of genome-wide divergence [1,4,7–10]. Thus, the challenge is to distinguish those targets of selection and demographic processes that generate the genomic landscape of divergence.

Here, we reconstruct the history of demographic isolation and characterize the extent to which selection has shaped genomic divergence between two closely related, hybridizing *Heliconius* species. More precisely, we first use a demographic modeling approach to reconstruct the history of population sizes and isolation during divergence of these species [11–18]. Next, with knowledge of the most likely demographic history, we use coalescent simulations to test for the importance of selection as the underlying mechanism driving heterogeneous patterns of genomic divergence. The coalescent simulation approach allows us to explore other genomic factors, such as recombination rate, on genomic divergence. Specifically, we can test for genome-wide impacts of linked selection by using the expectation that linked selection is higher in regions of the genome where recombination rate is lower [19]. Hence, we would expect a negative association between recombination rate and divergence across the genome [8,9,20,21]. Similarly, we expect a positive association between recombination rate and admixture, if the species continue to hybridize [4,22], but not necessarily if they diverged in isolation [8,23]. Thus, the relationships of recombination rate with divergence and admixture can be used to infer the relative importance of different evolutionary processes during speciation.

To provide a relative perspective of the divergence between our focal hybridizing *Heliconius* species, we first investigate the relationship between reproductive isolation and genomic divergence across 15 pairs of increasingly divergent populations and species in the *H. erato* clade. Next, we use the incipient species *Heliconius erato* and *Heliconius himera* that hybridize across three geographically distinct contact zones to test for the relative contribution of demographic and selective factors in the evolution of the divergence landscape. *Heliconius himera* is found in dry forest areas of southern Ecuador and northern Peru [24]. It comes into contact with *Heliconius erato cyrbia* on the western slopes of the Ecuadorian Andes and with *Heliconius erato favorinus* and *Heliconius erato emma* on the eastern slopes of the Andes, both areas with wet forest (Figure 1A). Hybridization is ongoing and hybrids are easily identifiable by their wing color patterns. In Ecuador, hybrids compose approximately 5% of the population in the contact zone [25], and, although poorly characterized, hybridization is known to occur in the other contact zones. The two species show strong premating isolation but little or no postmating reproductive barriers [25]. The eastern and western *H. erato* populations that hybridize with *H. himera* do not come into contact with each other and show deep genetic divergence in the *H. erato* clade [26].

**Figure 1.**
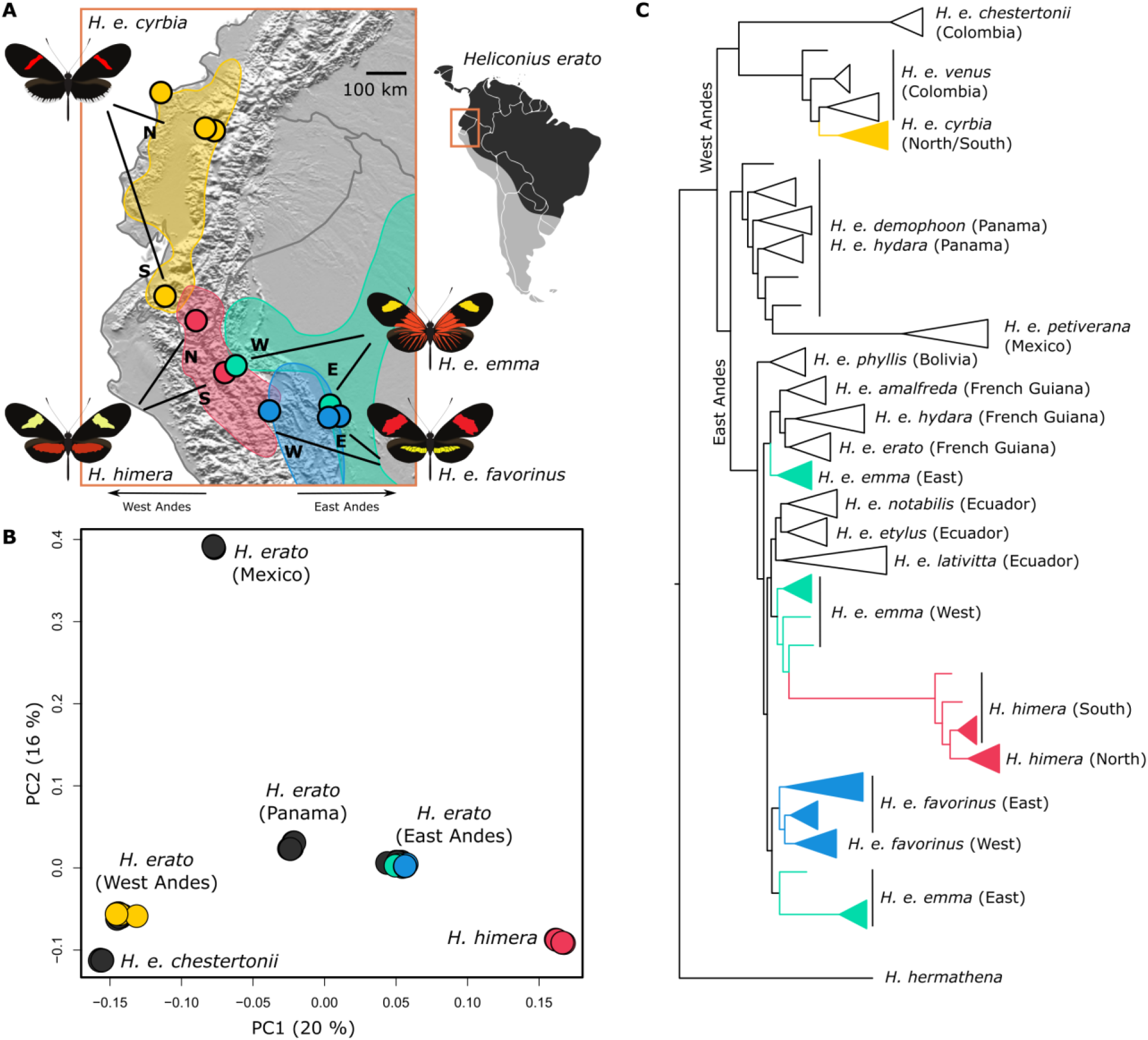
Geographical distribution, population structure and phylogeny of the focal populations in relation to the *Heliconius erato* radiation. **(A)** We sampled two populations of *H. himera*, *H. e. cyrbia*, *H. e. emma* and *H. e. favorinus*. The distribution of *H. himera* (red) covers dry valleys in the Andes of South Ecuador and North Peru. In the North, *H. himera* (N) comes into contact with a *H. e. cyrbia* (S) population. In the South, *H. himera* comes into contact with a *H. e. emma* (W) and *H. e. favorinus* (W) population. **(B)** Principal Component Analysis (PCA) of the focal samples (colored points) among all the available whole genome data for the *H. erato* radiation (black points). **(C)** Maximum likelihood tree built using FastTree and using only autosomal sites from 121 whole genome resequenced individuals (see Figure S1 for the uncollapsed tree). Nodes in the tree that represent the major clades within *H. erato* (east and west of Andes) obtained high support (= 1) from the Shimodaira-Hasegawa test.

Our findings demonstrate the importance of both isolation and selection in establishing the heterogeneous genomic landscape of divergence that characterizes our hybridizing species. The reconstructed demographic history supports a complex dynamics of population fluctuations and varying migration rates, that with selection, resulted in the observed heterogeneous patterns of genomic divergence. Further, our coalescent simulations well fit the observed relationship of genomic divergence and recombination when we consider genome-wide impacts of recent and strong selective events in the absence of gene flow. Finally, we show that the species boundary between *H. himera* and *H. erato* is highly porous and that gene flow is highly asymmetrical and distinct across each of the three contact zones. We suggest that this asymmetrical signal of gene flow may be the result of polygenic species boundaries that restrict introgression in *H. himera*. Overall, our results highlight that the study of heterogeneous landscapes of divergence can help us understand how demographic and selective processes drive speciation.

## Results & discussion

### Genomic landscape of divergence among hybridizing races and incipient species

We first sought to describe the general patterns of genome-wide divergence and how they varied based on the varying degrees of reproductive isolation. To do this, we compared genome-wide patterns of divergence across 15 contact zones in the *H. erato* clade that have varying degrees of reproductive isolation (*RI*, Figure 2; Table S1). Correlations of *RI* with genome-wide estimates of relative divergence (*F_ST_*) show that divergently selected color patterns between hybridizing races of *H. erato* with absence of other pre- or post-mating barriers are not sufficient to drive genome-wide increases in divergence (Figure 2A). In these hybridizing *H. erato* races, there are only narrow peaks of divergence largely centered over the loci known to be responsible for color pattern differences (Figure 2B). In contrast, between the incipient species *H. erato cyrbia* and *H. himera*, which have strong differences in mate preference, divergence is much higher across the entire genome. Divergence between these species has increased to the extent that *F_ST_* peaks near the known color pattern loci *WntA* (chr 10), *cortex* (chr 15), and *optix* (chr 18) are not detectable (Figure 2B). The *H. himera* and *H. erato* contact zones on the eastern Andes (*H. e. emma* and *H. e. favorinus*) also show elevated genome-wide divergence, but much lower overall than in the western Andes contact zone. As expected, genomic divergence was highest between *H. e. venus* and *H. e. chestertonii* from Colombia, where both mate preference and hybrid sterility have been reported [27]. We also note a dramatic increase in divergence on the Z chromosome relative to the autosomes (Figure 2). This is in line with previous work on *H. e. chestertonii*, which suggested the important role played by the Z chromosome as a barrier to gene flow [28].

**Figure 2.**
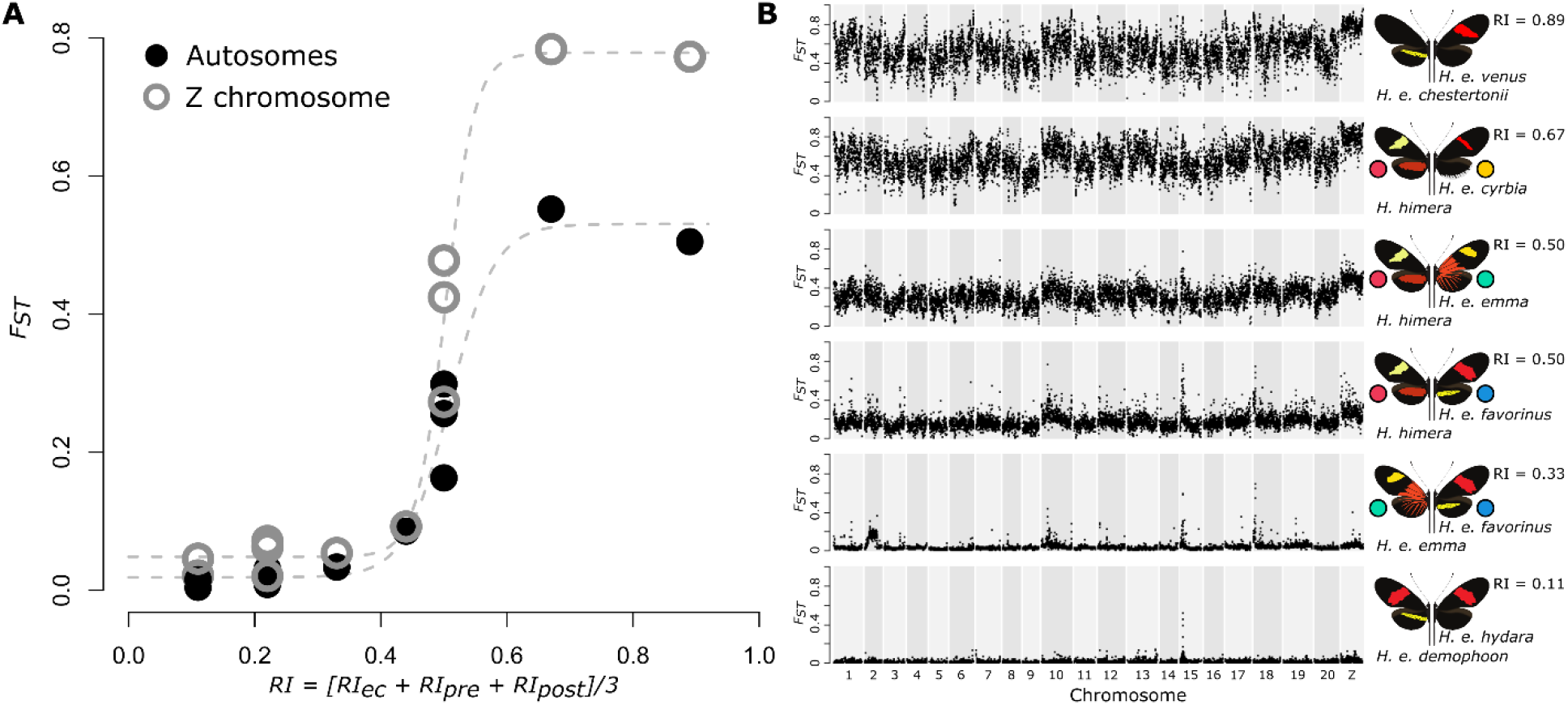
Reproductive isolation and divergence among *Heliconius erato* populations. **(A)** Genome-wide averages of relative divergence (*F_ST_*) show a sharp increase with increasing measures of reproductive isolation between parapatric *H. erato* populations. The measure of reproductive isolation (*RI*) was obtained by equally weighting ecological (*RI_ec_*), pre-isolation (*RI_pre_*) and post-isolation (*RI_post_*) components (Table S1). Higher relative divergence on the Z chromosome can be observed for the more divergent parapatric comparisons, however, incompatibilities that are potentially Z-linked have only been suggested for *H. e. chestertonii* crosses [27,28]. **(B)** Plots of relative divergence (average *F_ST_* in 50 kb windows) between parapatric *H. erato* populations along the genome. Plots are ordered according to the measure of reproductive isolation (*RI*). Colored circles match color codes used for the focal populations in this study. Divergence peaks on chromosome 10, 15 and 18 correspond to the divergently selected color pattern genes *WntA* (affecting forewing band shape), *cortex* (affecting yellow hindwing bar), and *optix* (affecting red color pattern elements), respectively [26].

### Demographic change and selection jointly drive genomic divergence among incipient species

Collectively, our results provide a view into the genomic landscape among lineages with increasing degrees of reproductive isolation (Figure 2)[29,30]. However, the increase in divergence does not seem to be a linear process as there is a marked increase in divergence between the incipient species that are known to still frequently and continuously hybridize over many generations. To understand what drives these elevated patterns of divergence, we have to recognize that each of these contact zones reflect hybridization between evolutionary distinct lineages. In this regard, the genomic landscapes do not reflect a continuum of genomic divergence throughout speciation, but rather they each are the evolutionary outcomes of various neutral and adaptive processes that shaped each of the populations coming into contact. For example, we find increased divergence between *H. himera* and *H. erato* from the western Andes slopes compared to *H. erato* from eastern Andes slopes. As seen in the PCA and phylogenetic inference, this results from a deeper split between *H. himera* and *H. e. cyrbia* from the western Andes slope compared to *H. himera* and the *H. erato* races that are found east of the Andes, including *H. e. emma* and *H. e. favorinus* (Figure 1B, C). This is consistent with previous studies that placed *H. himera* nested within the *H. erato* clade and not as a sister species to *H. erato* [26,31].

To understand what forces are likely driving differences in patterns of genome-wide divergence between *H. erato and H. himera*, we fit the estimated joint site-frequency spectrum (JSFS) for three geographically distinct contact zones to 26 alternative demographic scenarios that varied in split times, migration rates and population sizes (Figure 3). All three *H. erato* and *H. himera* contact zones best fit models that included secondary contact (SC) after a period of isolation without gene flow (Figure S2-3; Table S2). For all three zones we found support for asymmetrical migration. In each case, migration rates were predominantly in one direction, with on average 0.5 to 0.6 migrants per generation moving from *H. erato* into *H. himera*, compared to 0.07 to 0.13 moving from *H. himera* into *H. erato*. This result is consistent with the effective migration rates being driven by the marked population size differences between the two species (Figure 3B).

**Figure 3.**
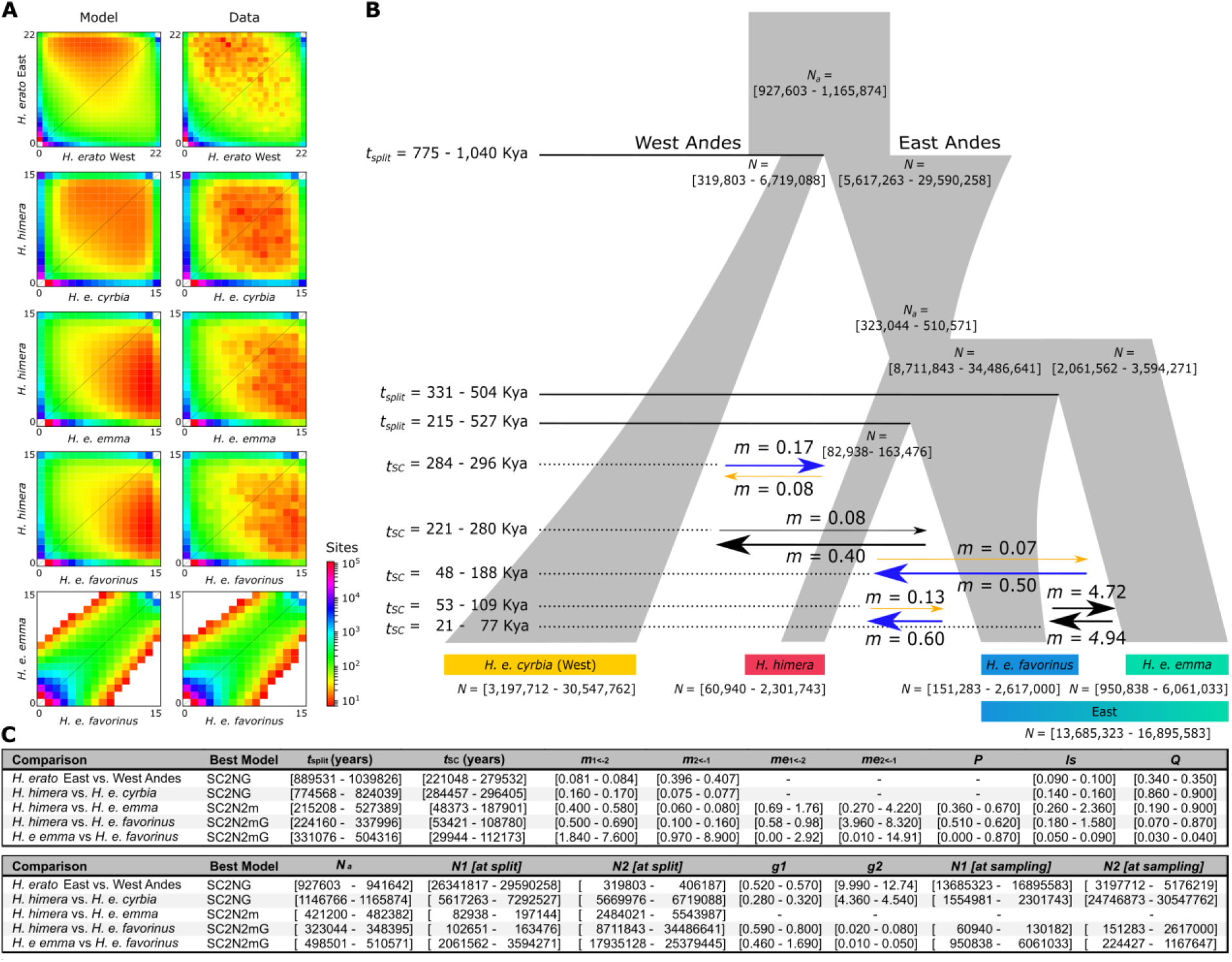
Secondary Contact (SC) best demographic model of the *H. himera* and *H. erato* population history. **(A)** Joint Site Frequency Spectra (JSFS) for data and best model (see Table S2 and Figure S3-4 for AIC values (Akaike Information Criterion)). **(B)** Reconstruction of historical demography of *H. himera* and *H. erato* populations using models with best AIC scores. All best models included a period of isolation and secondary contact. Arrows indicate effective migration rates (2*N_a_m*). Migration from *H. himera* into *H. erato* is indicated in orange, migration from *H. erato* into *H. himera* is indicated in blue. **(C)** Table with parameter ranges obtained from five best scoring models out of twenty runs. *N_a_* = ancestral population size, *N_1_* = Size of population 1, *N_2_* = size of population 2, *g*1 = growth coefficient of population 1, *g*2 = growth coefficient of population 2, *ls* = linked selection, *Q* = proportion of the genome with a reduced effective size due to linked selection (*ls*), *m*_1<−2_ = migration from population 2 into 1, *m*_2<−1_ = migration from populations 1 into 2, *t*_split_ = split time, *t*_SC_ = time of secondary contact, *P* = proportion of the genome evolving neutrally.

Nearly all the models that included exponential population growth (G) best fit the JSFS, with the exception of the *H. himera* and *H. e. emma* comparison. Estimates of ancestral and contemporary population sizes suggest strong expansions in *H. himera* and *H. erato*. These inferred changes in population sizes broadly fit previous results obtained from pairwise sequentially Markovian coalescent (PSMC) analysis (Van Belleghem et al. 2018), which suggested an overall population growth in *H. erato* east and west of the Andes in the past 1 My and size reduction for *H. himera* in the past 200 Ky (Figure S3). However, we found that estimates of contemporary population sizes varied greatly depending on the population comparison (Figure 3B, C), a result possibly explained by unaccounted population structure and difficulties in estimating growth (G). For *H. e. favorinus,* estimates of contemporary population size were generally much smaller than the ancestral population. This result fits the observation that *H. e. favorinus* is a smaller Andean population of the “postman” color pattern (i.e. red forewing band), which has a much larger distribution throughout the Neotropics. Collectively, the models support a history that includes periods of allopatry, followed by lineage specific changes in population size that coincide with more recent gene flow.

To investigate if the JSFS contained evidence of selection driving patterns of divergence across the contact zones, we incorporated heterogeneity in population size (2N) and migration rate (2M) into the models, similar to what was done by Rougeux *et al.* 2017 [32] and Tine *et al.* 2014 [16], respectively. The 2N model allows heterogeneity in population size estimates across loci that result from the differences in allelic variation caused by linked selection (*ls* = effective population size of locus relative to neutral loci; Q = proportion of the genome affected by *ls*). We found that all contact zones between *H. erato* and *H. himera* well supported 2N models, suggesting the effect of linked selection in shaping patterns of genomic variation and divergence between the incipient species. The strongest *ls* was observed for the population comparisons of *H. erato* East and West (*ls* = 0.10; *Q* = 0.35) and *H. himera* and *H. erato* West (*ls* = 0.15; *Q* = 0.90) and the lowest observed between *H. e. emma* and *H. e. favorinus* (*ls* = 0.07; *Q* = 0.04) (Figure 3C, Table S3).

The result of selection on locally adapted alleles is a heterogeneous landscape with regions containing these variants showing much lower rates of admixture compared to the rest of the genome. The 2M models allow for this type of heterogeneity in migration rates across the genome. Both contact zones in the eastern Andes supported these models for *H. himera* and *H. erato*, suggesting that the model fits the eastern Andean populations having genomic regions with much lower rates of introgression than other parts of the genome.

### Simulations of linked selection and secondary contact predict correlation of recombination rate with divergence, not admixture

The demographic models provide a comprehensive reconstruction of the evolutionary history of divergence between *H. erato* and *H. himera* and allow us to estimate time intervals of lineage splits, size changes and migration changes that span the past million year of divergence between *H. himera* and *H. erato* in the Andes (Figure 3). We next use these estimates to inform simulations and conduct further tests for the impact of selection and demography during genomic divergence of the incipient species. To do this, we simulated the expected relationship of recombination rate to relative divergence (*F_ST_*) and admixture (*f_d_*) at a locus linked to a site under selection. We used two distinct scenarios of migration broadly applicable *to H. erato* and *H. himera* and compared the simulation results to real data (Figure 4A, B).

**Figure 4.**
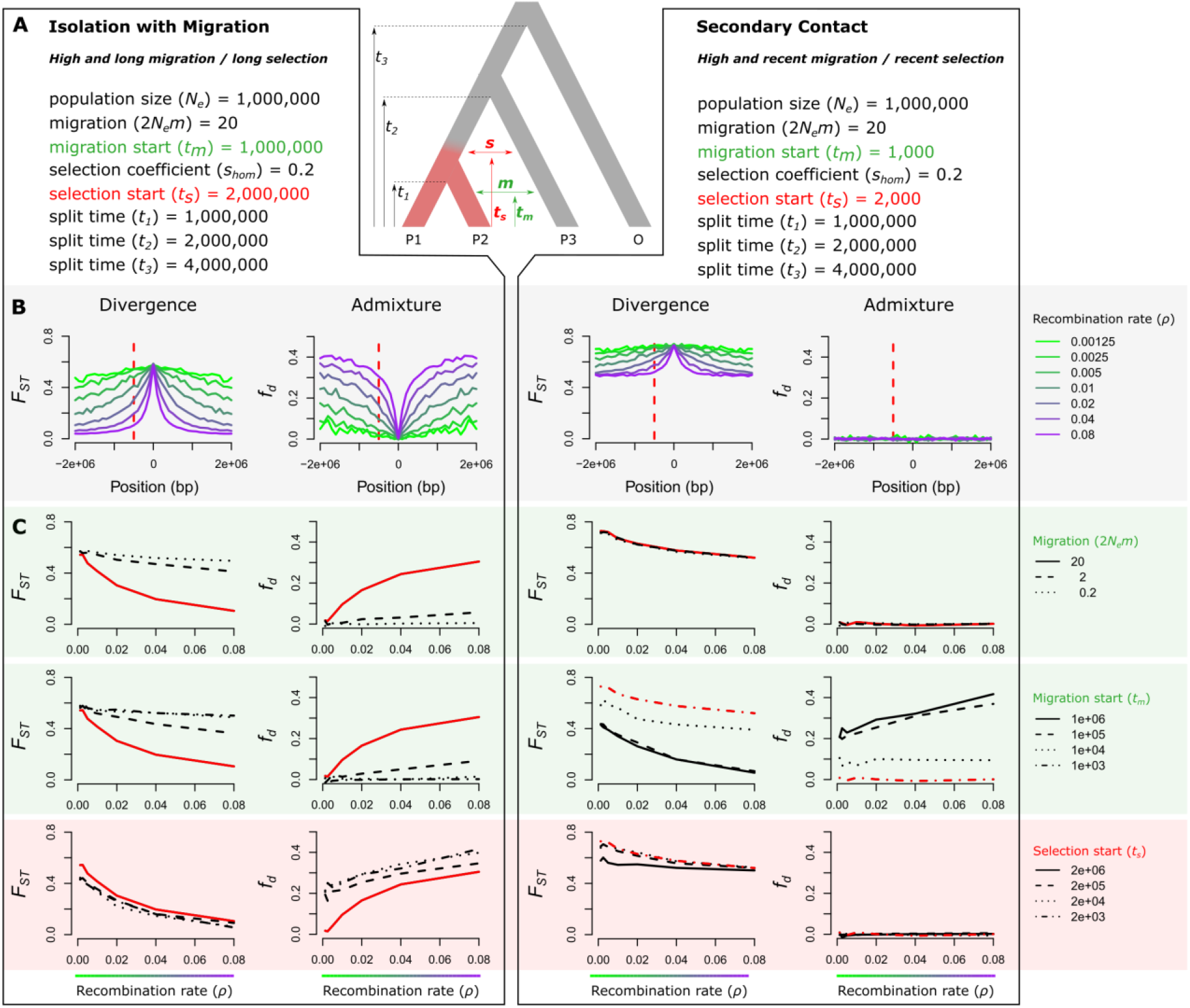
Expected relationship of recombination rate with divergence (*F_ST_*) and admixture (*f_d_*) near a divergently selected locus. **(A)** The population tree shows the simulated scenario with the onset of divergent selection on a derived allele indicated in red and in which migration rate (*m*) and migration time (*t_m_*) between P2 and P3 and selection start time (*t_s_*) are varied. Left and right of the simulated scenario are parameter combinations for two extreme scenarios that both include linked selection; on the left a scenario with Isolation with Migration (IM) and on the right a scenario reflecting Secondary Contact (SC). **(B)** Effect of population recombination rate (*ρ*) on relative divergence (*F_ST_*) and admixture (*f_d_*) near a divergently selected locus for the two simulated scenarios with parameter combinations as in panel A. The selected allele occurs at position 0. The dashed red line indicates a locus at 500 kb from the selected locus at which the relationship between *ρ*, divergence and admixture is assessed in panel C. **(C)** The effect of migration start time (*t_m_*) and selection start time (*t_s_*) on the relation between *ρ*, divergence and admixture. Apart from the respective parameters being evaluated, other parameters were fixed as in panel A, with the red lines indicating the exact parameter combinations as in panel A and B. For simulations with a lower selection coefficient (*s* = 0.02), see Figure S5.

As demonstrated by other studies, in a scenario of “isolation with migration” (IM) with long periods of migration and divergent selection, our simulations predict a strong negative correlation between recombination rate and *F_ST_* [20,21,33] and a strong positive correlation between recombination rate and *f_d_* [4,8] (Figure 4C, left). This pattern reflects the degree of linkage between neutral sites and loci under divergent selection [23,34]. In contrast, our simulations show that the relationship between recombination rate and *f_d_* is reduced when migration is more recent or low (Figure 4C, left). As expected, in a “secondary contact” (SC) scenario that is characterized by a more recent onset of migration, the simulations show that *F_ST_* values are generally high and *f_d_* values close to zero (Figure 4C, right). This results in the absence of a relationship between recombination rate and *F_ST_* or *f_d_* in most SC scenarios. However, when selection in the genome is recent (~selective sweeps), a negative relationship between recombination rate and divergence, but not admixture, emerges in the SC scenario (Figure 4C). This relationship arises due to linked selection that reduces diversity within populations and increases the relative divergence, *F_ST_*, between populations [35]. The relationship holds for lower selection strengths at relatively short distances from the selected site (s = 0.02; Figure S5).

### Patterns of divergence are shaped by linked selection

The differences in the expected relationships between recombination rate and *F_ST_* and between recombination rate and *f_d_* under the IM and SC scenarios provide specific predictions that we can use to determine how linked selection may have shaped genomic divergence between *H. erato* and *H. himera*. We first explored this relationship of recombination rate, relative divergence and admixture across the *optix* locus using the expected patterns obtained from our simulations. The *optix* locus controls red wing pattern differences among *H. erato* races, as well as between *H. erato* and *H. himera* and is a target of strong selection (Figure 5A) [26,31,36]. We found recombination rate (*ρ*) estimates were markedly lower upstream, compared to downstream of the *optix* gene (Figure 5C). Such sharp decreases in recombination rate near chromosome ends have been observed in other *Heliconius* species [4] and likely explain the decrease in recombination rate upstream of the *optix* gene. To test if selection at the *optix* locus has produced the expected relationship of recombination rate with divergence and admixture, as predicted above in our simulations (Figure 4C), we sampled sites 500 kb from the center of the peak of the divergence at *optix* and plotted the recombination rate and divergence estimates for those sites (Figure 5D). As expected from the scenario of SC with linked selection, we found negative relationships between recombination rate and divergence (*F_ST_*) and no correlations between recombination rate and admixture (*f_d_*). These findings demonstrate that our simulated patterns of recombination, divergence and admixture, reflect real observed patterns in a region known to be under strong selection.

**Figure 5.**
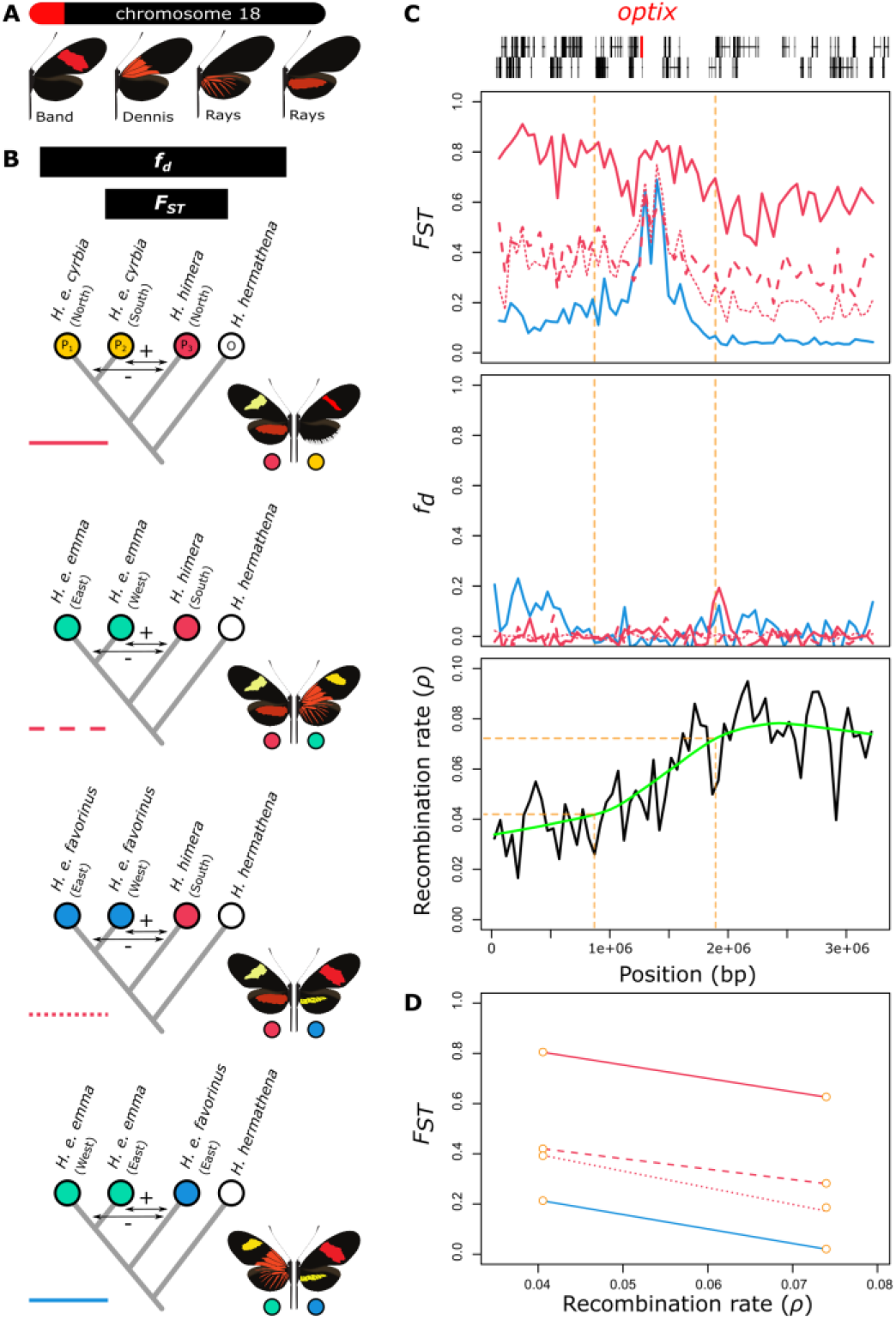
Divergence (*F_ST_*), admixture (*f_d_*) and recombination rate (*ρ*) near the red color pattern gene *optix*. **(A)** The *optix* gene is located near the start of chromosome 18 and has been demonstrated to control the expression of red color pattern elements in *Heliconius* wings [37]. **(B)** Relative divergence (*F_ST_*) and admixture (*f_d_*) comparisons performed between *H. himera*, *H. e. cyrbia*, *H. e. emma* and *H. e. favorinus*. Colored circles match color codes in Figure 2. **(C)** Lines show *F_ST_*, *f_d_* and recombination rate (*ρ*) calculated in 50 kb non-overlapping windows. The green line in the bottom plot shows a loess fit of the recombination rate. Gene models including the location of the *optix* gene are presented at the top. **(D)** Relationship between *F_ST_* and recombination rate (*ρ*) 500 kb left and right from the center of the *optix* regulatory sequence divergence peak.

To test for evidence that linked selection has driven genome-wide patterns of divergence, we compared recombination rates with divergence and admixture across the whole genome. We found a significant negative association between recombination rate (*ρ*) and relative divergence (*F_ST_*) between *H. himera* and *H. erato* populations (Figure 6A) but no association with admixture (*f_d_*) (Figure 6B). These genome-wide patterns are identical to those observed at the *optix* locus (Figure 5D) and the simulations of SC with linked selection (Figure 4C). Further, we observed a positive association between proportion of coding sequence and relative divergence, which again suggests the importance of linked selection for genome divergence (Figure S6). Additionally, we found a significant relationship between *F*_*ST*_ and *f*_*d*_ in the *H. himera* – *H. erato* hybrid zones, which suggests that, although admixture can partly explain reduced *F_ST_* (Figure 6C), the rates of migration have been too low or recent to build up a significant association with recombination rate. Finally, we note that the observed patterns of linked selection may include the effects of both genetic hitchhiking and background selection. While our genomic dataset does not allow us to differentiate between them, our simulations suggest that the observed patterns can be explained by genetic hitchhiking alone and other studies suggest background selection may be too subtle to cause these patterns [9].

**Figure 6.**
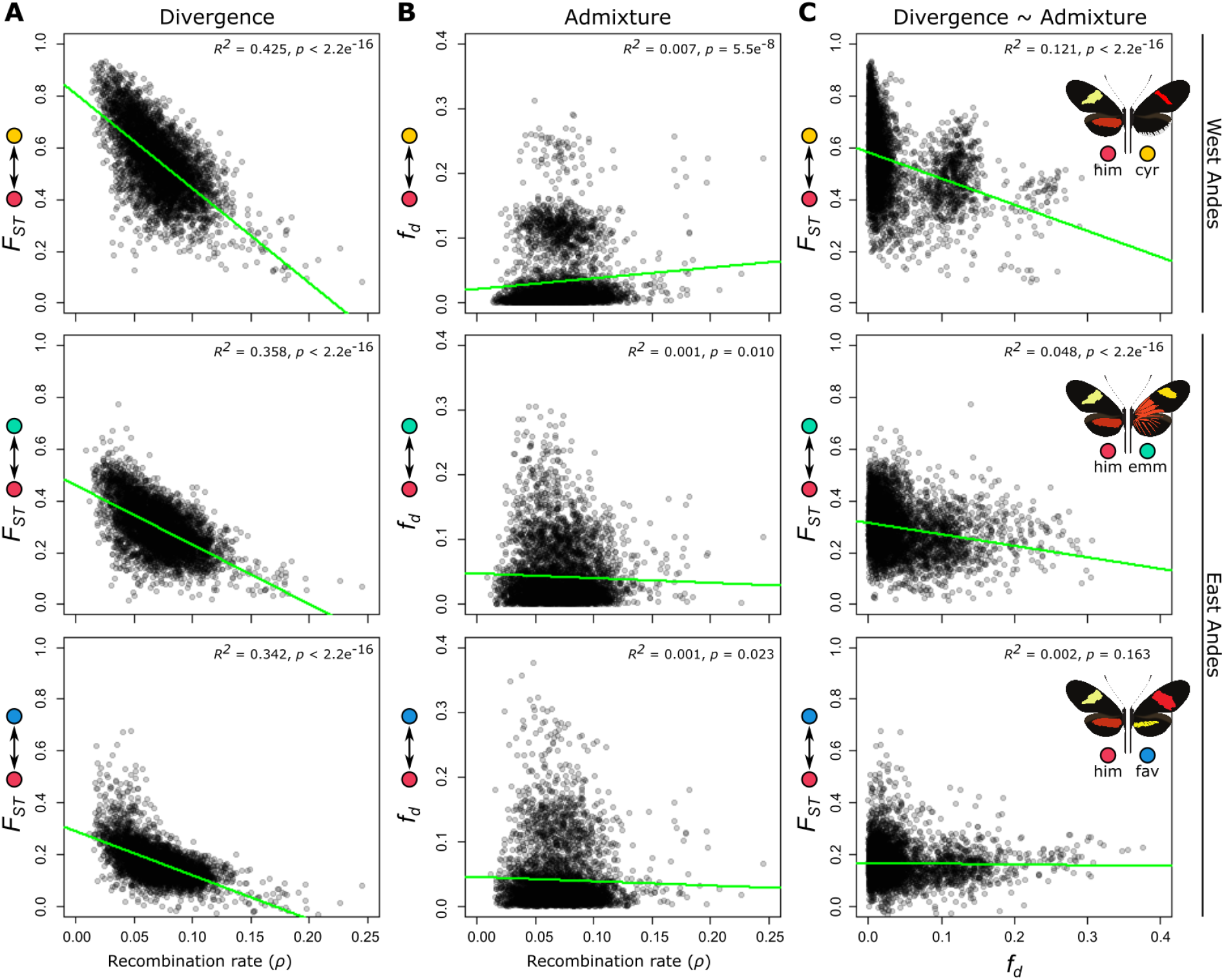
Correlations of divergence and admixture proportions with recombination rate in the three *H. himera – H. erato* contact zones. **(A)** Relative divergence (*F_ST_*) versus recombination rate (*ρ*). **(B)** Admixture proportion (*f_d_*) versus recombination rate (cM/Mb). **(C)** Relative divergence (*F_ST_*) versus admixture proportion (*f_d_*). Statistics were calculated in 50 kb non-overlapping windows. Recombination rates were calculated from each *H. erato* population separately and averaged over populations (see methods) and showed a genome-wide average of *ρ* equal to 0.071 (SD = 0.026; *ρ* = 4*N*_e_*r*). Colored circles match geographic distributions and contact zones in Figure 2.

### Asymmetrical admixture suggests a polygenic species boundary

The combination of empirical and simulation data suggests that periods of isolation and linked selection within populations have played a major role in shaping the divergence landscape between *H. erato* and *H. himera*. To explore how these factors influence the finer details of genomic divergence we investigated patterns of admixture along chromosomes. We were both interested in the patterns of admixture across our three replicate hybrid zones and inferring the direction of admixture. To determine if patterns of admixture were similar across the three contact zones, we used a modified four-taxon *D*-statistic called *f_d_* [38]. Across several chromosomes we observed large blocks of increased *f_d_*, particularly in the hybrid zone between *H. himera* and *H. e. cyrbia* west of the Andes (see chromosomes 3, 4, 6, 7, 8, 12 and 18 in Figure 7A). The large size of these admixture tracks indicates that these signals are recent and there has not been sufficient time for recombination to break down the haplotypes. In contrast, high admixture values (*f_d_*) between *H. himera* and *H. e. emma* and *H. himera* and *H. e. favorinus* east of the Andes are distributed more evenly along the chromosomes (Figure 7A), which would agree with older admixture events between these populations. In general, there was a lack of overlap between genomic regions with high *f_d_* in the different hybrid zone comparisons further reinforcing the idea that we are examining independent admixture events.

**Figure 7.**
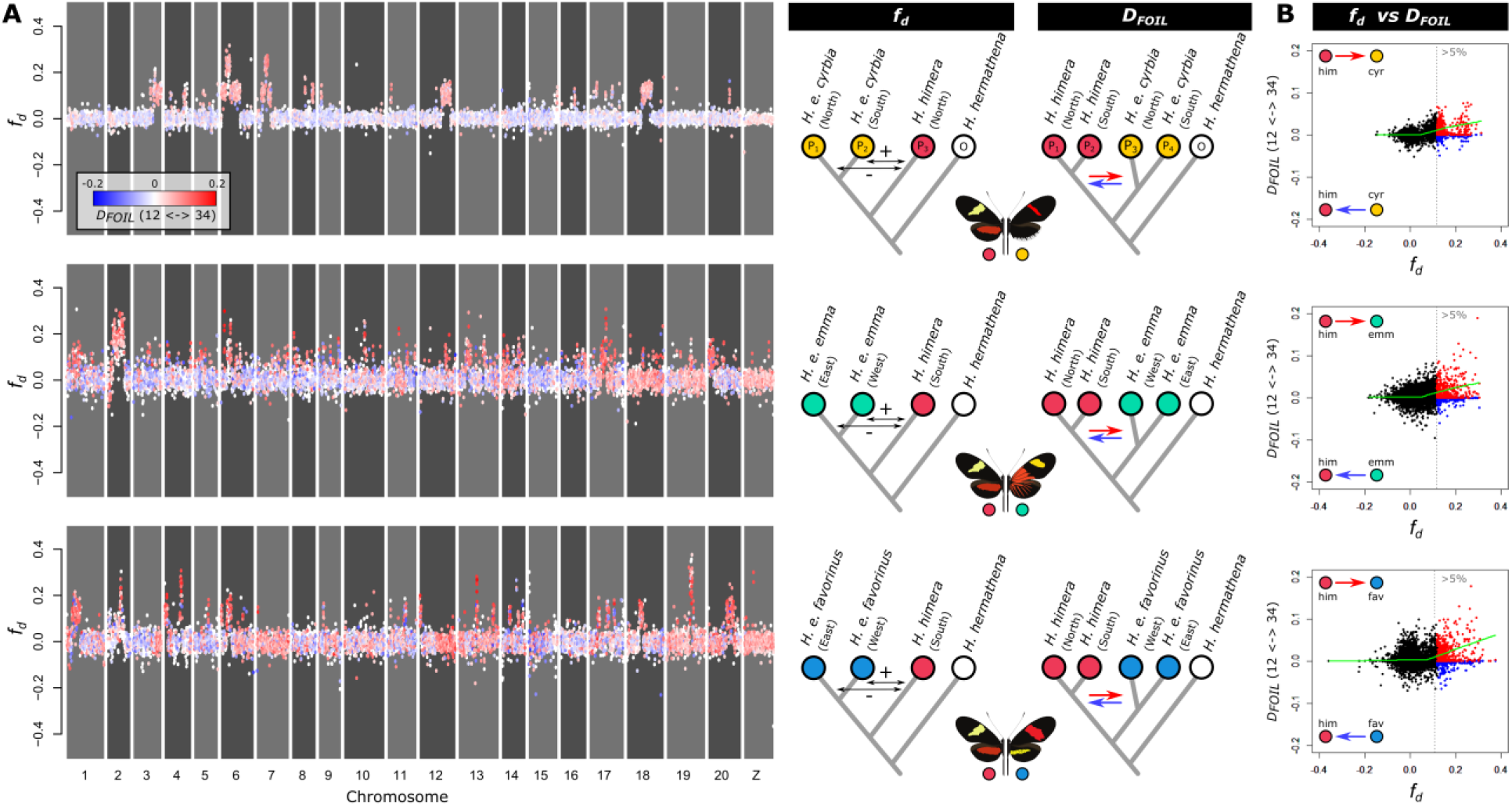
Admixture (*f_d_*) and admixture directionality (*D_FOIL_*) between *H. himera* and *H. erato* in three contact zones. **(A)** Points show admixture (*f_d_*) values, whereas coloring shows directionality (*D_FOIL_*) in 50 kb non-overlapping windows for the contact zones *H. himera* – *H. e. cyrbia* (top), *H. himera* – *H. e. emma* (middle) and *H. himera* – *H. e. favorinus* (bottom). Blue indicates predominant admixture from *H. erato* into *H. himera* (12 <− 34), whereas red indicates predominant admixture from *H. himera* into *H. erato* (12 -> 34) based on the *D_FOIL_* tests. **(B)** Summary of admixture versus directionality with points above the 95% quantile indicated in blue (12 <− 34) and red (12 -> 34) demonstrates that the majority of windows with high *f_d_* indicate admixture from *H. himera* into *H. erato*. The green line indicates a loess fit of the data. Colored circles match color codes in Figure 2.

To determine the directionality of introgression across the genome we used a five-taxon *D_FOIL_*-statistic [39]. This test considers all available taxa (i.e. two populations of *H. himera* and two populations of *H. erato*) and calculates all possible four-taxon *D-*statistic comparisons to infer admixture as well as directionality of admixture. While the *D_FOIL_*-statistic is calculated using a single genome for each taxon, we performed this test on all possible combinations of available samples and represented the *D_FOIL_* signal as the proportion of significant comparisons (Figure 7A). Comparing *f_d_* and *D_FOIL_* results revealed that among loci that show strong evidence of admixture, there is a relative paucity of loci in *H. himera* individuals carrying *H. erato* alleles (Figure 7B). This asymmetry suggests *H. erato* is more porous to introgressed alleles from *H. himera* than vice-versa. Consistent with a well characterized “large X(Z) effect” in speciation, on the sex (Z) chromosome there are only a few loci that show signals of admixture (high *f_d_*) and again the *D_FOIL_* tests only show evidence of introgression of *H. himera* alleles into *H. erato* (Figure 7A).

We propose that the multitude of loci with asymmetrical gene flow may represent the genetic signal of a polygenic species barrier. These barriers could result from co-adapted loci in *H. himera* that cannot be replaced by *H. erato* alleles without fitness consequences. The inference that this pattern results from a polygenic species barrier is further strengthened by two observations. First, *H. himera* males have been shown to mate more frequently with F1 hybrids compared to *H. erato* males [25], which should result in an opposite asymmetric admixture pattern than we found [40], with more alleles introgressing from *H. erato* into *H. himera*. Second, the smaller effective population size estimates of *H. himera* should also result in an opposite pattern of asymmetric admixture [13], with more alleles introgressing from *H. erato* into *H. himera.* This latter expectation is observed in the demographic modeling results, which show greater migration of alleles from *H. erato* into *H. himera* and corresponds to their estimated differences in population size between the species (Figure 3B), but is not seen in the more recent signals of admixture as measured by *f_d_*.

### Different histories can generate similar heterogeneous divergence patterns

Although the genomic landscape of divergence between hybridizing taxa reflects the history of selection and demographic changes they have experienced, different histories can generate strikingly similar heterogeneous patterns. This can greatly mimit our ability to make inferences about the evolutionary processes driving genomic change [2]. For example, in *Heliconius melpomene* there are strikingly similar heterogeneous landscapes of divergence to those we report here for *H. erato* and *H. himera*, despite known differences in their evolutionary histories [41,42]. Here, we show that through a comprehensive set of analytical approaches we can disentangle these evolutionary histories and reveal that different evolutionary processes have resulted in similar divergence landscapes among the *H. erato* and *H. melpomene* clades.

*Heliconius melpomene* and *H. erato* are co-mimics and experience similar strong selective pressures on wing color patterns throughout their distributions. In the *H. melpomene* clade, *H. melpomene* comes into contact and hybridizes with *H. cydno* in Panama and occasionally with *H. timareta* in Ecuador and northern Peru. Although at first sight the genomic landscapes appear similar, correlation analyses reveal differences in the relative importance of admixture. The *H. melpomene* comparisons showed strong correlations of recombination rate with divergence as well as admixture, which supports a longer history of divergent selection with gene flow (Martin et al. 2019). In contrast, *H. himera* and *H. erato* comparisons showed recombination rate correlated with divergence, but not admixture. We suggest the lack of correlation may be because the secondary contact is recent and the rates of gene flow are low between *H. erato* and *H. himera*. Thus, there has not been enough time for a correlation between recombination rate and admixture to arise. Instead, we argue that differences accumulated in periods of isolation have had more profound effects on the divergence landscape of *H. erato* and *H. himera* whereas gene flow has likely been more continuous throughout the divergence history of the young species pair in the *H. melpomene* clade. These differences in the history of divergence among co-mimetic species highlights the power of approaches like those applied here to resolve the roles of different evolutionary processes in generating seemingly similar heterogeneous patterns of genomic divergence.

### Conclusions

We observed genome-wide increase of divergence as reproductive barriers increase between hybridizing populations or species. Nonetheless, it remains difficult to determine if peaks of divergence result from barrier loci that are resistant to ongoing gene flow (heterogeneous gene flow), from recent selective sweeps in isolated populations (heterogeneous selection), or both. Fortunately, using a combination of demographic modeling, simulations, and correlation analyses we can characterize the evolutionary processes responsible for the heterogeneous landscapes of divergence. For the incipient species *H. erato* and *H. himera*, our results suggest a disproportionately large effect of the Z chromosome on the evolution of species barriers in *H. erato*. The data also favor a scenario of periods of isolation accompanied by both selection and gene flow. The overall patterns of asymmetric admixture suggest that during periods of isolation, selection at multiple loci may have resulted in a polygenic species boundary. This finding adds to a number of studies that illustrate how fluctuating gene flow and pervasive selection along the genome lead to the evolution of polygenic architectures of species boundaries [4,8,21,43,44].

## Methods

### Sampling

We obtained whole genome resequence data for a total of 122 *Heliconius* butterflies (Tables S4). These include northern (North Ecuador, n = 10) and southern (South Ecuador *H. himera* contact zone, n = 4) *H. e. cyrbia*, western (Peru *H. himera* contact zone, n = 4) and eastern (Peru *H. e. favorinus* contact zone, n = 7) *H. e. emma* and western (Peru *H. himera* contact zone, n = 4) and eastern (Peru *H. e. emma* contact zone, n = 8) *H. e. favorinus* used to study admixture patterns with northern (Ecuador *H. e. emma/favorinus* contact zone, n = 5) and southern *H. himera* (Peru *H. e. cyrbia* contact zone, n = 4). Additionally, samples from *H. e.* petiverana (Mexico, n = 5), *H. e. demophoon* (Panama, n = 10), *H. e. hydara* (Panama, n = 6 and French Guiana, n = 5), *H. e. erato* (French Guiana, n = 6), *H. e. amalfreda* (Suriname, n = 5), *H. e. notabilis* (Ecuador *Heliconius erato lativitta* contact zone, n = 5 and Ecuador *H. e. etylus* contact zone, n = 5), *H. e. etylus* (Ecuador, n = 5), *H. e. lativitta* (Ecuador, n = 5), *H. e. phyllis* (Bolivia, n = 4), *H. e. venus* (Colombia, n = 5) and *H. e. chestertonii* (Colombia, n = 7) were used for contact zone divergence analysis and population structure visualization as well as samples of *H. hermathena* (Brazil, n = 3) as an outgroup to root the phylogenetic inference and polarize site frequency spectra. All data have been previously published [26,28], apart from the ten *H. e. cyrbia* from North Ecuador.

### Sequencing and genotyping

Genotypes were obtained as in [28]. In short, whole-genome 100-bp paired-end Illumina resequencing data from *H. erato* samples were aligned to the v1 [26] reference genome, using BWA v0.7 [45]. PCR duplicated reads were removed using PICARD v1.138 (http://picard.sourceforge.net) and sorted using SAMTOOLS [46]. Genotypes were called using the genome analysis tool kit (GATK) Haplotypecaller [47]. Individual genomic VCF records (gVCF) were jointly genotyped using GATK’s genotype GVCFs. Genotype calls were only considered in downstream analysis if they had a minimum depth (DP) ≥ 10, maximum depth (DP) ≤ 100 (to avoid false SNPs due to mapping in repetitive regions), and for variant calls, a minimum genotype quality (GQ) ≥ 30. Specific data filtering steps for running population structure, phylogenetic and demographic analysis are explained in the respective sections.

### Population structure and phylogenetic relationships

We estimated levels of relative divergence (*F_ST_*) [48] between populations in nonoverlapping 50 kb windows using python scripts and egglib [49]. For this analysis, we only considered windows for which at least 10% of the positions were genotyped for at least 75% of the individuals within each population. On average 96% of windows met this criterium. To discern population structure among the *H. erato* and individuals, we performed principal component analysis (PCA) using EIGENSTRAT SmartPCA [50]. For this analysis, we only considered autosomal biallelic sites that had coverage in all individuals and excluding the Z chromosome (5,058,785 SNPs). Using the same filtering but including the outgroup *H. hermathena* (4,927,152 SNPs), we used FastTree v2.1 [51] to infer an approximate maximum likelihood phylogeny using the default parameters.

### Demographic modeling

We performed demographic analyses on the joint site-frequency spectra (JSFS) of five population comparisons using a modified version of *∂a∂i* v1.7 [11]. Genotype calls were filtered for biallelic autosomal SNPs with a threshold of at least 50 % minimum genotype calls for each population of interest. In order to ensure demographic analyses were performed on unlinked loci, we subsampled our data so that a single SNP was selected at least every ~1000 bp (based on linkage disequilibrium maps from [52] in Figure S1). Unfolded joint site-frequency spectra (JSFS) were created from the filtered calls data using *H. hermathena* as an outgroup, including on average 316090.3 SNPS (Table S3). The proportion of accurate SNP orientation (O) was consistent across all pairwise comparisons (~97 %), which suggests that ancestral state identification was correct for the majority of SNPs using *H. hermathena* as an outgroup.

Models tested include four basic scenarios in addition to 22 extensions of the basic models that allowed for independent assessment of additional selective and demographic parameters [32] (Table S5). Basic model scenarios included strict isolation (SI), ancient migration (AM), isolation-with-migration (IM), and secondary contact (SC). The standard four models involved the divergence of an ancestral population (*N_ref_*) at *t* generations into two resulting daughter populations with an effective size of *N_1_* and *N_2_*, respectively. Migration occurred in IM, AM, and SC at rate *m_12_* from population 2 into population 1 and *m_21_* in the opposite direction. Model extensions included growth rate parameters (*g*) which account for changes in the effective population size over time (bottlenecks and expansions) by taking a ratio of the contemporary and ancient population sizes. Note that *∂a∂i* cannot infer both heterogeneous migration and genetic drift when gene flow is not temporally localized (i.e. IM model). To capture the effects of linked selection that are due to sweeps and background selection (Hill and Robertson, 1966; Maynard Smith and Haigh, 1974; Charlesworth et al, 1993), further model extensions incorporated two categories of loci (2*N*) that occur in proportions of *Q* and 1-*Q* in order to account for heterogeneity in *N_e_* across the genome. A Hill-Robertson scaling factor (*hrf*) was included with these models that relates the effective size of loci experiencing selection to that of neutral loci. Models that infer migration (IM, AM, SC) were extended to include parameters that capture heterogeneous migration rates (2*m*) across the genome resulting from selection on barrier loci during adaptive divergence [16]. These extensions allowed for the estimation of a proportion of loci (*P*) experiencing standard migration rates (*m_12_* and *m_21_*) and a second category of loci (1-*P*) undergoing rates *me_12_* and *me_21_*. Given that the JSFS was polarized using *H. hermathena* as the outgroup for all comparisons, a SNP orientation parameter (*O*) was included in all models to account for ancestral state misidentification. Additionally, the effective mutation rate (*ϑ*) of *N_ref_* was estimated as a free parameter in all comparisons.

To check for model convergence, a total of 20 independent optimizations were performed for each model on each population comparison. When running these models, consistency in the likelihood scores generally increased as the best optimized parameters from previous steps were incorporated into subsequent steps. To score the models and account for overparameterization, the Akaike Information Criterion (AIC: Burnham and Anderson 2004) was used and parameters from the top five scoring runs for each model were averaged. The models with the best average AIC score for each comparison were retained for each comparison. The highest and lowest optimized parameter values in the top five replicate runs for each model were used to construct intervals to estimate uncertainty.

Model parameter estimates for effective population sizes, migration rates and divergence times were transformed into absolute units using a *Heliconius* mutation rate of 2 × 10^−9^ per generation (i.e. spontaneous *Heliconius* mutation rate corrected for selective constraint; Keightley et al. 2014; Martin, Eriksson, et al. 2015) and assuming a generation time of 0.25 years. Ancestral effective population size was calculated using the optimized theta value for each model comparison (*N_ref_ = ϑ/4μL*), where L represents the effective sequence length and *μ* the estimate of the mutation rate. Effective sequence length for each pairwise comparison is estimated as *L = (x/y)z*, where *x* is the number of SNPs used in the ∂a∂i demographic analysis and *y* is the number of segregating sites detected in the original sample out of *z* total sites. Migration rates are presented in units of *2N_ref_* to represent the number of individuals per generation that migrate into each population.

### Divergence and admixture simulations

To compare patterns in our data to expectations, we simulated genealogies near a selected locus. Genealogies were simulated using the coalescent simulator *msms* [56] and from these genealogies 10 kb sequences were simulated with a mutation rate of 2e^−9^ (i.e. spontaneous *Heliconius* mutation rate corrected for selective constraint; Keightley et al. 2014; Martin, Eriksson, et al. 2015) and a Hasegawa–Kishino–Yano (HKY) substitution model using *seq-gen* v1.3.4 [57]. With *msms*, we simulated four populations with a split history given by (((P1, P2), P3), O) where *t_1_*, *t_2_* and *t_3_* denote the split times between P1 and P2, (P1, P2) and P3 and ((P1, P2), P3) and O, respectively (Figure 4A). Population size (*N_e_*) was fixed to 1,000,000 individuals and *t_2_* and *t_3_* were fixed to 0.5 (2,000,000 generations) and 1 (4,000,000 generations) coalescent units (4*N_e_*), respectively. Within the range of selection on color pattern in *Heliconius* [58], we simulated divergent selection at a single locus with selection coefficients (*s*) of 0.2 and 0.02 for homozygous genotypes and 0.1 and 0.01 for heterozygous genotypes and selection strength specified in units of 2*N_e_s*. Selection was set to work in opposite direction for P1 and P2 versus P3 and O and was set to start at time *t_s_*. After population splits, migration (*m*) was restricted between P2 and P3 only, with symmetrical migration rates and a start time equal to *t_m_*. In relevance to our demographic modeling results, we ran simulations by varying the parameters *t_s_* (selection start time; 2e^3^ - 2e^6^ generations), *t_m_* (migration start time; 2.5e^−5^ - 5e^−1^ generations), *m* (migration rate; 1e^−7^ - 1e^−5^ generations) and *ρ* (population recombination rate 4*N_e_r*; *r* = probability of recombination per generation per bp; 6.25e^−4^ - 8e^−2^). A maximum *ρ* of 8e^−2^ was used for computational feasibility. The genealogies were sampled at 100 kb increments from the selected locus. This was achieved by using an infinite recombination sites model and changing the position of the selected locus in increments of 10 neutral locus units (i.e. 10 × 10 kb) away from the sampled locus. Divergence (*F_ST_*) was calculated as in Hudson *et al.* 1992 and admixture (*f_d_*) was calculated as in Martin *et al.* 2016 using python scripts and *egglib* v3 (De Mita & Siol, 2012). We investigated the correlation between *F_ST_*, *f_d_* and recombination rate at a distance of 500 kb from a selected locus but similar expectations are obtained from wide range of distances to the selected locus. Simulations were run with 100 replicates for each parameter combination. Pseudocode to run the *msms* command lines are provided in Tables S6.

### Recombination rate estimates

We estimated fine-scale variation in population recombination rate (*ρ* = 4*N*_e_*r*; *r* = probability of recombination per generation per bp) along the *H. erato* chromosomes from linkage-disequilibrium in population genetic data using *LDhelmet* v1.7 [59]. We phased quality filtered genotypes from thirteen *H. erato* populations (i.e. *H. e. cyrbia*, *H. e. venus*, *H. e. demophoon*, *H. e. hydara* (Panama), *H. e. emma*, *H. e. etylus*, *H. e. lativitta*, *H. e. notabilis*, *H. e. favorinus*, *H. e. phyllis*, *H. e. erato*, *H. e. hydara* (French Guiana) and *H. e. amalfreda*) using Beagle v4.1 [60] with default parameters. From the phased genotypes, fasta sequences were generated for 50 kb windows. These 50 kb windows were transformed to haplotype configuration files with the recommended window size of 50 SNPs used by *LDhelmet* to estimate composite likelihoods of the recombination rate. From the haplotype configuration files, lookup tables for two-locus pairwise recombination likelihoods and Padé coefficients were generated within the recommended value range. Transition matrices were calculated for each chromosome separately by comparing genotypes obtained from *H. erato demophoon* to the outgroup species *H. hermathena*. The likelihood lookup tables, Padé coefficients and transition matrices were used in the rjMCMC procedure of *LDhelmet* to estimate the recombination map. In this latter step, 1000,000 Markov chain iterations were run with a burn-in of 100,000 iterations, a window size of 50 SNPs and block penalty of 50. To reduce the potential effect of locus-specific changes in effective population size (*N_e_*) on population recombination rate (*ρ*) estimates (e.g. due to population specific selective sweeps or background selection), we estimated *ρ* for each *H. erato* population separately and obtained averages for each 50 kb interval.

### Admixture statistics

We estimated admixture proportions for 50 kb non-overlapping windows using the *f_d_* statistic [38]. This statistic is based on the ABBA-BABA test or Patterson’s *D* statistic which measures an excess of derived allele sharing between sympatric non-sister taxa [61]. This excess is tested by comparing the relative abundance of SNP patterns termed ABBAs and BABAs in a tree of three populations and an outgroup with the relationship (((P1, P2), P3), O). ABBAs are sites where a derived allele is shared between P2 and P3, whereas BABAs are sites where a derived allele is shared between P1 and P3. Under a neutral coalescent model, such sites are only expected to be found due to incomplete lineage sorting or recurrent mutation and a *D* statistic of 0 is expected. In the presence of admixture, however, an access of either ABBAs or BABAs can be observed and a D statistic that significantly deviates from 0 may be obtained. The *f*_d_ statistic is derived from the *D* statistic by calculating the difference between ABBA and BABA sites and normalizing this difference by a scenario of complete admixture. The estimator is dynamic in that for the complete admixture scenario used to normalize, a donor population for the admixture is chosen with the highest frequency of the derived site. The resulting normalized measure is approximately proportional to the effective migration rate and has been evaluated not to be confounded by locus-specific changes in effective population size due to background selection or reductions in diversity due to selective sweeps [4,38]. The populations included as P1, P2, P3 and O are indicated in the figures. *Heliconius hermathena* samples were consistently used as the outgroup taxa.

### Admixture directionality

By expanding the four-taxon *D* statistic to a five-taxon scenario, it is possible to obtain information on the directionality of admixture (i.e. donor versus recipient population). A set of statistical measures that use a five-taxon symmetric phylogeny to infer both the taxa involved in and the direction of admixture are called the *D_FOIL_* statistics [39]. The *D_FOIL_* statistics identify taxa involved in admixture by performing four possible *D* tests with different combinations of three ingroup taxa within a five-taxon phylogeny defined as (((P1, P2), (P3, P4)), O). These four *D* tests considered collectively can provide information on the directionality of admixture. This is because admixture does not only change the position of the donor sample in the topology but will also change the relationship of the donor’s sister taxon to the other taxa in the phylogeny. For instance, if admixture occurs from P2 into P3, the sampled topology becomes (((P2, P3), P1), P4), O) and P1 will group more closely to (P2, P3) because of more recent sharing of variation with P2, whereas if admixture occurs from P3 into P2, the topology becomes (((P2, P3), P4), P1), O). For the latter instance, this will be reflected by a similar sign of the *D* test statistics that include (((P1, P2), P3), O) or (((P1, P2), P4), O) but a different sign for the D test that includes (((P3, P4), P2), O) and no significant D test for (((P3, P4), P1), O). Hence, by comparing the combinations of different signs (+, −, or 0) of the four *D* tests within the five-taxon topology, the directionality of admixture can be assessed [39].

We assessed directionality of admixture in 50 kb non-overlapping windows in the three *H. himera* contact zones with *H. erato* populations using the *D_FOIL_* tests explained above using the available *dfoil* software (www.github.com/jbpease/dfoil). Samples from the *H. himera* North and South populations were specified as the P1 and P2 group, whereas samples from the considered *H. erato* populations were specified as P3 and P4. *Heliconius hermathena* was used as the outgroup taxon (ID hermathena_13 in Table S4). The *D_FOIL_* statistics were calculated between each possible combination of available ingroup taxa (i.e. one sample for each taxon group); 800 combinations for *H. himera* – *H. e. cyrbia*, 500 for *H. himera* – *H. e. emma* and 480 for *H. himera* – *H. e. favorinus*. Among these sample combinations, significant *D_FOIL_* signatures (χ^2^ goodness-of-fit test) were counted and used to obtain heterogeneous patterns of admixture directionality along the genome.

## Data accessibility

For GenBank accession numbers of whole genome resequence data see Table S4.

## Author contributions

The study was conceived and designed by SVB and BAC in collaboration with RP and WOM. Genomic analyses were performed by SVB and JC. Demographic modeling with *∂a∂I* was conducted by JC. Samples of *H. e. cyrbia* from northern Ecuador were contributed by GMK, and CB assisted with permits. RP, GMK and WOM provided input on results and manuscript preparation. The manuscript was written and figures were made by SVB, JC, and BAC.

## Acknowledgments

This work was funded by NSF grant (DEB 1257689) to BAC and RP and NSF EPSCoR RII Track-2 FEC (OIA 1736026) to RP and BAC. For sequencing and computational resources, we thank the University of Puerto Rico, the Puerto Rico INBRE Grant P20 GM103475 from the National Institute for General Medical Sciences (NIGMS), a component of the National Institutes of Health (NIH); and awards 1010094 and 1002410 from the EPSCoR program of the NSF. We thank the Ecuadorian Ministerio del Ambiente (No. 005-13 ICFAU-DNB/MA), Peruvian Ministerio de Agricultura and Instituto Nacional de Recursos Naturales (201-2013-MINAGRI-DGFFS/DGEFFSS) and Autoridad Nacional De Licencias Ambientales-ANLA in Colombia (Permiso Marco 0530) for permission to collect butterflies. We thank Simon Martin for sharing his code and useful discussions that shaped this paper. We thank Nicola Nadeau for access to sequence data for northern *H. erato cyrbia* samples, which was funded by her UK Natural Environment Research Council (NERC) fellowship (NE/K008498/1).

## Notes

### Competing Interest Statement

The authors have declared no competing interest.

