## Supplementary material for "Selection and gene flow define polygenic barriers between incipient butterfly species"

**Table S1. Hybrid zone characteristics between *H. erato* and *H. himera* populations.** Major color pattern (CP) loci include the genes *optix* (~red), *WntA* (~forewing band shape) and *cortex* (~yellow hindwing bar and white fringes) Reproductive isolation (*RI*) was estimated as a combination of ecological isolation ($RI_{ec}$) due to divergent selection on color patterns, pre-mating isolation ($RI_{pre}$) due to mate preference, and post-mating isolation ($RI_{post}$) due to hybrid sterility.

| race/species 1 | race/species 2 | Country | HZ width (km) | Major CP loci | RI pre | RI post | $RI_{ec}$ | $RI_{pre}$ | $RI_{post}$ | *RI* |
|---|---|---|---|---|---|---|---|---|---|---|
| *H. e. demophoon* | *H. e. hydara* | Panama | 50 | 1 | no | no | 0.33 | 0 | 0 | 0.11 |
| *H. e. erato* | *H. e. amalfreda* | Suriname | unknown | 1 | likely no | no | 0.33 | 0 | 0 | 0.11 |
| *H. e. hydara* | *H. e. amalfreda* | Suriname | 60 | 2 | likely no | no | 0.67 | 0 | 0 | 0.22 |
| *H. e. hydara* | *H. e. erato* | Suriname | 20 | 2 | likely no | no | 0.67 | 0 | 0 | 0.22 |
| *H. e. hydara* | *H. e. erato* | French Guiana | 20 | 2 | likely no | no | 0.67 | 0 | 0 | 0.22 |
| *H. e. favorinus* | *H. e. emma* | Peru | 10 | 3 | likely no | no | 1 | 0 | 0 | 0.33 |
| *H. e. notabilis* | *H. e. lativitta* | Ecuador | 20 | 2 | likely no | no | 0.67 | 0 | 0 | 0.22 |
| *H. e. notabilis* | *H. e. etylus* | Ecuador | unknown | 2 | likely no | no | 0.67 | 0 | 0 | 0.22 |
| *H. e. lativitta* | *H. e. etylus* | Ecuador | unknown | 1 | likely no | no | 0.33 | 0 | 0 | 0.11 |
| *H. e. favorinus* | *H. himera* | Peru | unknown | 3 | likely | no | 1 | 0.5 | 0 | 0.50 |
| *H. e. emma* | *H. himera* | Peru | unknown | 3 | likely | no | 1 | 0.5 | 0 | 0.50 |
| *H. e. etylus* | *H. himera* | Ecuador | unknown | 3 | likely | no | 1 | 0.5 | 0 | 0.50 |
| *H. e. cyrbia (south)* | *H. himera* | Ecuador | 5 | 3 | color based assortative mating | no | 1 | 1 | 0 | 0.67 |
| *H. e. venus* | *H. e. cyrbia (north)* | Colombia | unknown | 2 | likely no | asymmetrical hybrid female sterility | 0.33 | 0.5 | 0.5 | 0.44 |
| *H. e. venus* | *H. e. chestertonii* | Colombia | unknown | 2 | color based assortative mating | asymmetrical hybrid female sterility | 0.67 | 1 | 1 | 0.89 |

**Table S2. AIC weights dadi models.** AIC scores for each model are averages of the top 5 runs. Shaded cells indicate lowest AIC values.

| **Pop1** | **East** | **him** | **him** | **him** | **him** | **emm** | **emmE** | **emmW** | **himN** | **himS** | **himS** | **himS** |
|---|---|---|---|---|---|---|---|---|---|---|---|---|
| **Pop2** | West | cyr | East | emm | fav | fav | favE | favW | cyrS | cyrN | emmW | favW |
| **Model** | | | | | | | | | | | | |
| ***AM*** | 26919.26 | 23075.75 | 2796.91 | 2954.21 | 4989.54 | 6362.25 | 1315.94 | 839.68 | 13755.05 | 12922.72 | 1599.09 | 1373.76 |
| ***AM2m*** | 15449.69 | 18040.85 | 2328.94 | 2485.63 | 4399.40 | 5405.21 | 998.65 | 675.38 | 11372.29 | 9719.13 | 1130.01 | 942.11 |
| ***AM2mG*** | 7941.90 | 5630.27 | 2367.99 | 2867.41 | 3580.90 | 4740.54 | 985.43 | 635.86 | 1784.70 | 3083.26 | 1340.91 | 1035.81 |
| ***AM2N*** | 9459.37 | 11034.97 | 2343.83 | 5734.03 | 4968.65 | 6881.48 | 948.14 | 684.20 | 8003.61 | 6427.68 | 2589.45 | 1476.20 |
| ***AM2N2m*** | 28802.32 | 11955.19 | 2344.28 | 2540.11 | 4804.03 | 5478.89 | 990.67 | 681.38 | 8523.30 | 6701.09 | 1131.01 | 1067.97 |
| ***AM2N2mG*** | 20333.27 | 9248.34 | 2358.85 | 2518.52 | 3471.55 | 4961.82 | 981.71 | 658.87 | 3273.95 | 4019.61 | 1072.01 | 727.11 |
| ***AM2NG*** | 8568.71 | 11476.83 | 2346.49 | 2531.90 | 3659.59 | 4960.60 | 936.65 | 638.13 | 4120.42 | 4623.60 | 1106.48 | 674.98 |
| ***AMG*** | 14116.01 | 7106.34 | 2724.88 | 3057.07 | 4102.82 | 6108.61 | 1297.69 | 817.29 | 2956.11 | 4797.53 | 1556.57 | 1193.26 |
| ***IM*** | 26915.23 | 23104.73 | 2815.33 | 2972.06 | 5052.31 | 6137.89 | 1314.65 | 900.28 | 14454.92 | 13099.18 | 1595.42 | 1497.55 |
| ***IM2m*** | 15311.14 | 18043.88 | 2327.40 | 2491.71 | 4398.22 | 5699.64 | 1012.81 | 823.66 | 11739.95 | 9717.71 | 1132.34 | 992.51 |
| ***IM2mG*** | 6861.70 | 4856.30 | 2278.96 | 2450.25 | 3522.82 | 5193.79 | 984.58 | 733.95 | 1538.84 | 2921.75 | 1128.24 | 992.97 |
| ***IM2N*** | 9856.47 | 14564.19 | 2364.59 | 2597.64 | 5049.57 | 5401.50 | 965.74 | 809.06 | 9385.81 | 10579.96 | 1132.24 | 1395.67 |
| ***IM2NG*** | 6391.23 | 5244.28 | 2335.25 | 2526.60 | 4019.25 | 5439.45 | 961.36 | 771.38 | 2193.78 | 2653.48 | 1158.55 | 950.78 |
| ***IMG*** | 14067.60 | 7107.68 | 2758.19 | 3014.62 | 4153.04 | 6188.51 | 1296.72 | 857.60 | 2956.77 | 4796.28 | 1544.48 | 1483.87 |
| ***SC*** | 21545.03 | 22348.57 | 2819.50 | 2972.82 | 5053.95 | 6053.70 | 1286.60 | 901.32 | 13788.37 | 12911.50 | 1563.82 | 1501.14 |
| ***SC2m*** | 12364.36 | 18283.69 | 2326.15 | 2476.72 | 4393.80 | 5615.49 | 1012.15 | 831.73 | 12362.81 | 9827.77 | 1058.25 | 994.92 |
| ***SC2mG*** | 8051.03 | 4831.21 | 2481.64 | 3739.86 | 4425.83 | 5390.52 | 1056.65 | 798.11 | 3075.11 | 2461.28 | 2025.09 | 1336.13 |
| ***SC2N*** | 8805.14 | 10309.65 | 2369.01 | 2600.70 | 5058.78 | 5916.22 | 968.10 | 830.84 | 7883.62 | 6287.76 | 1143.50 | 1403.46 |
| ***SC2N2m*** | 9473.62 | 10668.07 | 2246.31 | 2428.11 | 4594.06 | 5696.28 | 981.90 | 855.23 | 7236.82 | 6496.57 | 1034.46 | 977.69 |
| ***SC2N2mG*** | 7255.99 | 4953.65 | 2319.34 | 2509.12 | 3337.41 | 4496.31 | 932.77 | 632.94 | 1229.20 | 2445.06 | 1029.17 | 832.20 |
| ***SC2NG*** | 6365.16 | 4610.33 | 2375.51 | 2641.85 | 3884.39 | 5029.93 | 957.76 | 718.66 | 1475.55 | 2548.38 | 1208.27 | 1058.73 |
| ***SCG*** | 13994.13 | 6890.99 | 2771.51 | 2975.06 | 4159.28 | 5756.15 | 1285.34 | 852.71 | 2822.26 | 4595.57 | 1516.51 | 1348.92 |
| ***SI*** | 58609.06 | 45115.69 | 20873.45 | 27209.19 | 33167.70 | 51478.88 | 23778.51 | 14467.52 | 27174.69 | 24736.58 | 15518.68 | 9581.81 |
| ***SI2N*** | 32469.12 | 32209.75 | 20267.44 | 25720.64 | 32879.09 | 51317.59 | 23640.42 | 14464.87 | 17982.67 | 7176.71 | 14723.36 | 9000.68 |
| ***SI2NG*** | 30005.12 | 18905.39 | 13120.81 | 18943.72 | 17924.01 | 44440.51 | 20150.49 | 11390.94 | 10241.78 | 7267.46 | 10751.43 | 5521.02 |
| ***SIG*** | 54201.47 | 25788.80 | 16559.54 | 23123.95 | 22442.43 | 45720.90 | 20261.97 | 11703.65 | 13094.69 | 11874.97 | 13989.10 | 7970.46 |

**Table S3. Best model parameters for best model class as well as alternative model classes.** Split times are given as Ttotal+TAM in AM, and Ttotal+TSC in SC. Although all best fitting demographic models include secondary contact (SC), several alternative models including AM (Ancient Migration) and IM (Isolation and Migration) fit relatively well to the observed JSFS (Figure 3A; Figure S2-3). However, in these models, migration rates generally scale proportionately with migration duration compared to the SC models, indicating that our interpretations on migration rates should not be affected by the uncertainty of finding a best demographic model.

| Pop1 | Pop2 | Best Model | AIC | log-likelihood | theta | theta/L |
|---|---|---|---|---|---|---|
| **East** | West | SC2NG | (6358.25, 6368.86) | (-3173.43, -3168.13) | (15445.43, 15679.19) | (0.0074, 0.0075) |
| **him** | cyr | SC2NG | (4586.15, 4691.20) | (-2334.60, -2271.39) | (42397.20, 43103.65) | (0.0092, 0.0093) |
| **him** | East | SC2N2m | (2205.15, 2302.05) | (-1139.02, -1090.57) | (10063.32, 11911.35) | (0.0042, 0.0050) |
| **him** | emm | SC2N2m | (2365.21, 2513.10) | (-1244.55, -1170.61) | (12981.08, 14866.68) | (0.0034, 0.0039) |
| **him** | fav | SC2N2mG | (3202.74, 3419.73) | (-1695.86, -1587.37) | (13234.76, 14273.34) | (0.0026, 0.0028) |
| **emm** | fav | SC2N2mG | (4305.65, 4618.41) | (-2295.21, -2138.83) | (11000.87, 11267.21) | (0.0040, 0.0041) |
| **East** | West | IM2NG | (6390.63, 6391.52) | (-3185.76, -3185.32) | (15644.64, 15728.41) | (0.0075, 0.0076) |
| **him** | cyr | IM2mG | (4852.69, 4860.10) | (-2419.05, -2415.35) | (47461.54, 49316.03) | (0.0103, 0.0107) |
| **him** | East | IM2mG | (2269.16, 2281.55) | (-1129.79, -1123.58) | (19496.36, 20340.74) | (0.0081, 0.0085) |
| **him** | emm | IM2mG | (2445.84, 2454.59) | (-1216.29, -1211.92) | (25565.14, 25716.54) | (0.0066, 0.0067) |
| **him** | fav | IM2mG | (3512.39, 3535.30) | (-1756.65, -1745.19) | (25506.39, 25877.43) | (0.0050, 0.0051) |
| **emm** | fav | IM2mG | (5094.93, 5323.25) | (-2650.63, -2536.47) | (21723.39, 23353.10) | (0.0079, 0.0085) |
| **East** | West | AM2mG | (7926.19, 7958.59) | (-3967.30, -3951.10) | (17512.82, 17870.57) | (0.0084, 0.0086) |
| **him** | cyr | AM2mG | (5581.90, 5666.58) | (-2821.29, -2778.95) | (37586.11, 45446.36) | (0.0081, 0.0098) |
| **him** | East | AM2m | (2328.84, 2329.22) | (-1154.61, -1154.42) | (20708.06, 20751.60) | (0.0086, 0.0087) |
| **him** | emm | AM2m | (2482.43, 2486.31) | (-1233.15, -1232.21) | (26200.25, 26226.96) | (0.00680, 0.00681) |
| **him** | fav | AM2N2mG | (3454.53, 3500.99) | (-1736.49, -1713.26) | (12740.53, 13038.79) | (0.0024, 0.0025) |
| **emm** | fav | AM2mG | (4694.64, 4835.32) | (-2405.66, -2335.32) | (21553.89, 22943.12) | (0.0078, 0.0083) |

| Pop1 | Pop2 | Nref | n1 | n2 | n1_final | n2_final |
|---|---|---|---|---|---|---|
| **East** | West | (927602.86, 941641.72) | (26341817.39, 29590257.54) | (319802.64, 406186.90) | (13685323.10, 16895583.18) | (3197711.72, 5176218.94) |
| **him** | cyr | (1146765.56, 1165873.74) | (5617263.04, 7292526.80) | (5669976.24, 6719087.99) | (1554980.53, 2301743.30) | (24746873.10, 30547761.60) |
| **him** | East | (525677.92, 622213.71) | (67830.57, 249820.76) | (2282101.19, 8015341.48) | - | - |
| **him** | emm | (421199.71, 482382.31) | (82937.57, 197143.88) | (2484020.60, 5543987.38) | - | - |
| **him** | fav | (323044.29, 348394.92) | (102651.12, 163475.82) | (8711842.83, 34486641.18) | (60940.22, 130181.67) | (151283.37, 2616999.91) |
| **emm** | fav | (498501.29, 510570.54) | (2061562.28, 3594270.56) | (17935127.69, 25379444.77) | (950838.31, 6061033.44) | (224427.38, 1167646.97) |
| **East** | West | (939566.71, 944597.44) | (6966144.6, 7086496.83) | (354416.53, 360027.67) | (30984262.05, 31901430.31) | (6441444.38, 6575968.28) |
| **him** | cyr | (1283746.70, 1333907.10) | (890091.34, 1034060.62) | (348415.79, 389006.23) | (509111.5, 646270.59) | (4368845.78, 5232016.85) |
| **him** | East | (1018432.06, 1062539.84) | (451063.98, 551350.03) | (4094835.02, 6224885.72) | (306964.56, 476021.17) | (8572225.92, 22252557.36) |
| **him** | emm | (829517.33, 834429.71) | (276050.86, 389567.33) | (4146735.54, 5139051.17) | (232565.46, 475272.85) | (8501981.32, 13915646.89) |
| **him** | fav | (622579.74, 631636.43) | (267012.48, 367516.17) | (265101477.72, 315168908.71) | (147568.75, 287135.21) | (1763171.64, 2571846.07) |
| **emm** | fav | (984389.56, 1058239.46) | (927371.18, 5568995.11) | (109352849.24, 622646834.2) | (1729621.1, 60334274.64) | (518111.04, 22801840.88) |
| **East** | West | (1051763.38, 1073248.76) | (8309755.63, 8653958.86) | (478065.91, 544040.23) | (16933042.70, 18413480.30) | (3890204.05, 5034447.94) |
| **him** | cyr | (1016634.71, 1229239.80) | (646807.81, 1065971.63) | (644077.54, 1046681.67) | (401442.26, 810581.39) | (3990167.40, 7346209.16) |
| **him** | East | (1081727.63, 1084002.08) | (338335.83, 340626.00) | (10660468.62, 10707372.65) | (233335.05, 234914.49) | (7352047.32, 7384394.93) |
| **him** | emm | (850124.99, 850991.62) | (294126.50, 295152.00) | (8621117.84, 8640601.67) | (202845.86, 203553.11) | (5945598.51, 5959035.64) |
| **him** | fav | (310980.88, 318260.85) | (44058.94, 70765.60) | (5656465.04, 7139282.62) | (1101707.65, 3416226.79) | (9905.86, 45475.89) |
| **emm** | fav | (976708.92, 1039660.72) | (1301099.94, 10066217.97) | (11055946.31, 102917154.38) | (955482.65, 197486732.69) | (19994.68, 5718393.61) |

| Pop1 | Pop2 | hrf | m12 | m21 | me12 | me21 |
|---|---|---|---|---|---|---|
| **East** | West | (0.09, 0.10) | (4.36E-08, 4.49E-08) | (2.14E-07, 2.16E-07) | - | - |
| **him** | cyr | (0.14, 0.16) | (6.79E-08, 7.30E-08) | (3.28E-08, 3.30E-08) | - | - |
| **him** | East | (0.30, 3.44) | (5.58E-07, 5.65E-07) | (5.96E-08 5.93E-08) | (5.56E-07, 3.74E-06) | (4.52E-07, 3.74E-06) |
| **him** | emm | (0.26, 2.36) | (4.73E-07, 6.04E-07) | (7.01E-08, 8.53E-08) | (8.23E-07, 4.37E-06) | (3.24E-07, 4.37E-06) |
| **him** | fav | (0.18, 1.58) | (7.74E-07, 9.91E-07) | (1.50E-07, 2.37E-07) | (8.91E-07, 1.20E-05) | (6.13E-06, 1.20E-05) |
| **emm** | fav | (0.05, 0.09) | (1.85E-06, 7.44E-06) | (9.72E-07, 8.71E-06) | (1.20E-09, 1.46E-05) | (6.95E-09, 1.46E-05) |
| **East** | West | (0.240, 0.242) | (2.827E-08, 2.828E-08) | (2.32E-07, 2.33E-07) | - | - |
| **him** | cyr | - | (3.43E-08, 3.45E-08) | (2.41E-08, 2.25E-08) | (7.82E-07, 8.06E-07) | (2.35E-07, 2.41E-07) |
| **him** | East | - | (2.1397E-07, 2.1399E-07) | (2.93E-08, 3.03E-08) | (4.38E-07, 8.89E-07) | (2.43E-07, 9.40E-07) |
| **him** | emm | - | (2.62E-07, 2.69E-07) | (3.80E-08, 4.11E-08) | (5.02E-07, 5.99E-07) | (1.08E-06, 1.22E-06) |
| **him** | fav | - | (3.42E-07, 3.67E-07) | (5.29E-08, 5.70E-08) | (1.60E-06, 1.98E-06) | (6.92E-07, 1.50E-06) |
| **emm** | fav | - | (5.30E-09, 7.40E-08) | (5.00E-10, 3.70E-07) | (2.88E-07, 1.04E-06) | (8.85E-07, 1.60E-06) |
| **East** | West | - | (6.32E-11, 4.61E-09) | (1.70E-07, 7.76E-08) | (6.48E-08, 6.89E-08) | (3.00E-07, 3.09E-07) |
| **him** | cyr | - | (4.92E-08, 4.15E-08) | (2.86E-09, 2.50E-10) | (9.97E-08, 9.98E-08) | (5.65E-08, 5.37E-08) |
| **him** | East | - | (9.67E-07, 9.85E-07) | (2.25E-07, 2.12E-07) | (2.22E-07, 2.24E-07) | (2.72E-08, 2.75E-08) |
| **him** | emm | - | (1.12E-06, 1.17E-06) | (2.81E-07, 2.66E-07) | (2.65E-07, 2.67E-07) | (3.77E-08, 3.80E-08) |
| **him** | fav | (0.10, 10.34) | (2.21E-06, 4.51E-06) | (2.22E-07, 1.31E-07) | (2.79E-07, 1.50E-05) | (7.26E-08, 1.44E-07) |
| **emm** | fav | - | (1.04E-12, 4.19E-08) | (4.05E-09, 4.94E-11) | (4.81E-08, 1.88E-06) | (8.62E-07, 1.80E-06) |

| Pop1 | Pop2 | Tsplit | Tam/Tsc | $T_{total}$ | P | Q | O |
|---|---|---|---|---|---|---|---|
| **East** | West | (668483.99, 760294.65) | (221047.09, 279531.79) | (889531.08, 1039826.44) | - | (0.34, 0.35) | (0.971, 0.972) |
| **him** | cyr | (490110.54, 527633.26) | (284457.48, 296405.25) | (774568.02, 824038.51 | - | (0.86, 0.90) | (0.966, 0.972) |
| **him** | East | (238584.88, 348057.41) | (66685.31, 223671.89) | (305270.19, 571729.30) | (0.37, 0.66) | (0.43, 0.90) | (0.97, 0.98) |
| **him** | emm | (166834.10, 339487.47) | (48373.42, 187901.15) | (215207.52, 527388.62) | (0.36, 0.67) | (0.19, 0.90) | (0.969, 0.975) |
| **him** | fav | (170738.66, 229216.34) | (53421.33, 108780.04) | (224159.99, 337996.37) | (0.51, 0.62) | (0.07, 0.87) | (0.973, 0.976) |
| **emm** | fav | (301132.78, 392142.62) | (29943.71, 112173.83) | (331076.49, 504316.45) | (0.00. 0.87) | (0.03, 0.04) | (0.970, 0.971) |
| **East** | West | (948190.31, 962353.23) | - | (948190.31, 962353.23) | - | (0.548, 0.544) | (0.9711, 0.9712) |
| **him** | cyr | (732629.31, 819439.32) | - | (732629.31, 819439.32) | (0.89, 0.90) | - | (0.968, 0.969) |
| **him** | East | (703822.40, 935721.10) | - | (703822.40, 935721.10) | (0.75, 0.77) | - | (0.968, 0.972) |
| **him** | emm | (689874.52, 720010.81) | - | (689874.52, 720010.81) | (0.79, 0.81) | - | (0.972, 0.973) |
| **him** | fav | (468698.71, 502136.70) | - | (468698.71, 502136.70) | (0.83, 0.87) | - | (0.9668, 0.9673) |
| **emm** | fav | (716006.23, 942620.30) | - | (716006.23, 942620.30) | (0.02, 0.23) | - | (0.970, 0.971) |
| **East** | West | (525037.07, 536574.75) | (259054.71, 292832.37) | (784091.78, 829407.12) | (0.36, 0.40) | - | (0.9725, 0.9728) |
| **him** | cyr | (378962.40, 622961.20) | (306675.48, 532410.46) | (685637.88, 1155371.66) | (0.34, 0.36) | - | (0.96, 0.97) |
| **him** | East | (705217.99, 709378.46) | (1.22, 281.19) | (705219.21, 709659.65) | (0.216, 0.224) | - | (0.9688, 0.9689) |
| **him** | emm | (589883.83, 590912.66) | (10.11, 229.99) | (589893.93, 591142.65) | (0.187, 0.195) | - | (0.9692, 0.9696) |
| **him** | fav | (227758.34, 248726.24) | (621.17, 2712.96) | (228379.51, 251439.19) | (0, 0.72) | (0, 0.08) | (0.9662, 0.9668) |
| **emm** | fav | (632115.02, 769059.75) | (13140.57, 195550.62) | (645255.59, 964610.37) | (0.01, 0.04) | - | (0.9698, 0.9703) |

**Table S4. Samples used in this study.** Samples from *H. himera* populations and *H. erato* populations that come into contact with *H. himera* are indicated in color. *H. himera* North, light red; *H. himera* South, dark red; *H. e. cyrbia* North, light yellow, *H. e. cyrbia* South, dark yellow; *H. e. emma* East, light green; *H. e. emma* West dark green; *H. e. favorinus* East, light blue; *H. e. favorinus* West, dark blue. Earthcape IDs are available through https://heliconius.ecdb.io.

| SequenceID | EarthCapeID | Taxon name | Country | Sex | Longitude | Latitude | NCBI Accession |
|---|---|---|---|---|---|---|---|
| BC2115 | BC2115 | *Heliconius erato amalfreda* | Suriname | m | -4.946897 | 55.183386 | SAMN05224103 |
| BC2124 | BC2124 | *Heliconius erato amalfreda* | Suriname | m | -5.943486 | 55.186072 | SAMN05224104 |
| STRI_WOM_5779 | STRI_WOM_5779 | *Heliconius erato amalfreda* | Suriname | m | -5.940653 | 55.189922 | SAMN05224208 |
| STRI_WOM_5780 | STRI_WOM_5780 | *Heliconius erato amalfreda* | Suriname | f | -5.940653 | 55.189922 | SAMN05224209 |
| STRI_WOM_5781 | STRI_WOM_5781 | *Heliconius erato amalfreda* | Suriname | f | -4.932733 | 55.200803 | SAMN05224210 |
| STRI_WOM_0057 | STRI_WOM_0057 | *Heliconius erato chestertonii* | Colombia | m | 3.884017 | -76.589367 | SAMN05224192 |
| STRI_WOM_0058 | STRI_WOM_0058 | *Heliconius erato chestertonii* | Colombia | f | 3.884017 | -76.589367 | SAMN05224193 |
| STRI_WOM_0059 | STRI_WOM_0059 | *Heliconius erato chestertonii* | Colombia | m | 3.884017 | -76.589367 | SAMN05224194 |
| 3661 | CS003661 | *Heliconius erato chestertonii* | Colombia | m | 3.884017 | -76.589367 | SAMN05224096 |
| 3662 | CS003662 | *Heliconius erato chestertonii* | Colombia | m | 3.884017 | -76.589367 | SAMN05224097 |
| 3663 | CS003663 | *Heliconius erato chestertonii* | Colombia | m | 3.884017 | -76.589367 | SAMN05224098 |
| 3664 | CS003664 | *Heliconius erato chestertonii* | Colombia | m | 3.884017 | -76.589367 | SAMN05224099 |
| cyrbia_004 | CYR004 | *Heliconius erato cyrbia* | Ecuador (S) | f | -3.726389 | -79.836667 | SAMN05224122 |
| cyrbia_005 | CYR005 | *Heliconius erato cyrbia* | Ecuador (S) | f | -3.726389 | -79.836667 | SAMN05224123 |
| cyrbia_023 | CYR023 | *Heliconius erato cyrbia* | Ecuador (S) | m | -3.726389 | -79.836667 | SAMN05224124 |
| cyrbia_024 | CYR024 | *Heliconius erato cyrbia* | Ecuador (S) | m | -3.726389 | -79.836667 | SAMN05224125 |
| CAM040673 | CAM040673 | *Heliconius erato cyrbia* | Ecuador (N) | m | 0.20981 | -78.95385 | SAMEA6447111 |
| CAM040695 | CAM040695 | *Heliconius erato cyrbia* | Ecuador (N) | m | 0.20981 | -78.95385 | SAMEA6447114 |
| CAM040861 | CAM040861 | *Heliconius erato cyrbia* | Ecuador (N) | m | 0.85369 | -79.80416 | SAMEA6447128 |
| CAM040853 | CAM040853 | *Heliconius erato cyrbia* | Ecuador (N) | m | 0.85369 | -79.80416 | SAMEA6447124 |
| CAM040578 | CAM040578 | *Heliconius erato cyrbia* | Ecuador (N) | m | 0.16174 | -78.75771 | SAMEA6447108 |
| CAM040692 | CAM040692 | *Heliconius erato cyrbia* | Ecuador (N) | m | 0.20981 | -78.95385 | SAMEA6447113 |
| CAM040584 | CAM040584 | *Heliconius erato cyrbia* | Ecuador (N) | m | 0.1641 | -78.75854 | SAMEA6447109 |
| CAM040842 | CAM040842 | *Heliconius erato cyrbia* | Ecuador (N) | m | 0.85369 | -79.80416 | SAMEA6447121 |
| CAM040545 | CAM040545 | *Heliconius erato cyrbia* | Ecuador (N) | m | 0.15135 | -78.76978 | SAMEA6447017 |
| CAM040860 | CAM040860 | *Heliconius erato cyrbia* | Ecuador (N) | m | 0.85369 | -79.80416 | SAMEA6447127 |
| Pet_ED3 | Pet_ED3 | *Heliconius erato demophoon* | Panama | m | -9.129444 | 79.715278 | SAMN05224182 |
| Pet_ED4 | Pet_ED4 | *Heliconius erato demophoon* | Panama | m | -9.129444 | 79.715278 | SAMN05224183 |
| Pet_ED5 | Pet_ED5 | *Heliconius erato demophoon* | Panama | m | -9.129444 | 79.715278 | SAMN05224184 |
| Pet_ED6 | Pet_ED6 | *Heliconius erato demophoon* | Panama | m | -9.129444 | 79.715278 | SAMN05224185 |
| STRI_WOM_0033 | STRI_WOM_0033 | *Heliconius erato demophoon* | Panama | f | -9.1525 | 78.689722 | SAMN05224188 |
| STRI_WOM_0082 | STRI_WOM_0082 | *Heliconius erato demophoon* | Panama | f | -9.1525 | 78.689722 | SAMN05224195 |
| STRI_WOM_0087 | STRI_WOM_0087 | *Heliconius erato demophoon* | Panama | m | -9.1525 | 78.689722 | SAMN05224196 |
| STRIWOM1284 | STRI_WOM_1284 | *Heliconius erato demophoon* | Panama | m | -9.1525 | 78.689722 | SAMN05224198 |
| STRIWOM5353 | STRI_WOM_5353 | *Heliconius erato demophoon* | Panama | f | -9.1525 | 78.689722 | SAMN05224202 |
| STRIWOM5362 | STRI_WOM_5362 | *Heliconius erato demophoon* | Panama | f | -9.1525 | 78.689722 | SAMN05224203 |
| BC2563 | BC_2563 | *Heliconius erato emma* | Peru | f | -5.29499 | -78.381 | SAMN08049955 |
| BC2577 | BC_2577 | *Heliconius erato emma* | Peru | m | -5.29499 | -78.381 | SAMN08049956 |
| BC2578 | BC_2578 | *Heliconius erato emma* | Peru | m | -5.29499 | -78.381 | SAMN08049957 |
| BC2579 | BC_2579 | *Heliconius erato emma* | Peru | m | -5.29499 | -78.381 | SAMN08049958 |

| | | | | | | | |
|---|---|---|---|---|---|---|---|
| GS020redo | GS020 | *Heliconius erato emma* | Peru | m | -6.181944 | -76.247222 | SAMN05224127 |
| GS021redo | GS021 | *Heliconius erato emma* | Peru | f | -6.181944 | -76.247222 | SAMN05224128 |
| NCS_1671 | NCS1671 | *Heliconius erato emma* | Peru | m | -6.181944 | -76.247222 | SAMN05224154 |
| NCS_1672 | NCS1672 | *Heliconius erato emma* | Peru | m | -6.181944 | -76.247222 | SAMN05224155 |
| NCS_1673 | NCS1673 | *Heliconius erato emma* | Peru | m | -6.181944 | -76.247222 | SAMN05224156 |
| NCS_1674 | NCS1674 | *Heliconius erato emma* | Peru | m | -6.181944 | -76.247222 | SAMN05224157 |
| NCS_1675 | NCS1675 | *Heliconius erato emma* | Peru | m | -6.181944 | -76.247222 | SAMN05224158 |
| NCS_2005 | NCS2005 | *Heliconius erato erato* | French Guiana | m | -4.638611 | 52.301667 | SAMN05224160 |
| NCS_2012 | NCS2012 | *Heliconius erato erato* | French Guiana | f | -4.638611 | 52.301667 | SAMN05224161 |
| NCS_2020 | NCS2020 | *Heliconius erato erato* | French Guiana | m | -4.585 | 52.245556 | SAMN05224162 |
| NCS_2023 | NCS2023 | *Heliconius erato erato* | French Guiana | m | -4.638611 | 52.301667 | SAMN05224163 |
| NCS_2025 | NCS2025 | *Heliconius erato erato* | French Guiana | m | -4.585 | 52.245556 | SAMN05224164 |
| NCS_2556 | NCS2556 | *Heliconius erato erato* | French Guiana | m | -4.621944 | 52.376111 | SAMN05224174 |
| BC_3277 | BC_3277 | *Heliconius erato etylus* | Ecuador | m | -1.97786 | -78.00945 | SAMN05224110 |
| BC_3278 | BC_3278 | *Heliconius erato etylus* | Ecuador | f | -1.97786 | -78.00945 | SAMN05224111 |
| BC_3280 | BC_3280 | *Heliconius erato etylus* | Ecuador | f | -1.97786 | -78.00945 | SAMN05224112 |
| BC_3281 | BC_3281 | *Heliconius erato etylus* | Ecuador | m | -1.97786 | -78.00945 | SAMN05224113 |
| BC_3282 | BC_3282 | *Heliconius erato etylus* | Ecuador | f | -1.97786 | -78.00945 | SAMN05224114 |
| BC2635 | BC_2635 | *Heliconius erato favorinus* | Peru | f | -6.4174 | -77.44329 | SAMN08049959 |
| BC2637 | BC_2637 | *Heliconius erato favorinus* | Peru | m | -6.4174 | -77.44329 | SAMN08049960 |
| BC2638 | BC_2638 | *Heliconius erato favorinus* | Peru | m | -6.4174 | -77.44329 | SAMN08049961 |
| GS012redo | GS012 | *Heliconius erato favorinus* | Peru | m | -6.461389 | -76.341944 | SAMN05224126 |
| NCS_0471 | NCS0471 | *Heliconius erato favorinus* | Peru | m | -6.474167 | -76.010278 | SAMN05224148 |
| NCS_0473 | NCS0473 | *Heliconius erato favorinus* | Peru | m | -6.474167 | -76.010278 | SAMN05224149 |
| NCS_0476 | NCS0476 | *Heliconius erato favorinus* | Peru | m | -6.474167 | -76.010278 | SAMN05224150 |
| NCS_0478 | NCS0478 | *Heliconius erato favorinus* | Peru | f | -6.474167 | -76.010278 | SAMN05224151 |
| NCS_0479 | NCS0479 | *Heliconius erato favorinus* | Peru | m | -6.474167 | -76.010278 | SAMN05224152 |
| NCS_2554 | NCS2554 | *Heliconius erato favorinus* | Peru | f | -6.474167 | -76.010278 | SAMN05224172 |
| NCS_2555 | NCS2555 | *Heliconius erato favorinus* | Peru | m | -6.474167 | -76.010278 | SAMN05224173 |
| STRI_WOM_0042 | STRI_WOM_0042 | *Heliconius erato hydara* | Panama | f | -9.1525 | 78.689722 | SAMN05224191 |
| NCS_1179 | NCS1179 | *Heliconius erato hydara* | French Guiana | m | -4.703611 | 52.303611 | SAMN05224153 |
| NCS_1979 | NCS1979 | *Heliconius erato hydara* | French Guiana | m | -4.571667 | 52.223333 | SAMN05224159 |
| NCS_2080 | NCS2080 | *Heliconius erato hydara* | French Guiana | f | -4.607778 | 52.2725 | SAMN05224165 |
| NCS_2211 | NCS2211 | *Heliconius erato hydara* | French Guiana | m | -4.547222 | 52.170278 | SAMN05224166 |
| NCS_2217 | NCS2217 | *Heliconius erato hydara* | French Guiana | m | -4.544444 | 52.1525 | SAMN05224167 |
| STRI_WOM_0039 | STRI_WOM_0039 | *Heliconius erato hydara* | Panama | m | -9.1525 | 78.689722 | SAMN05224189 |
| STRI_WOM_0040 | STRI_WOM_0040 | *Heliconius erato hydara* | Panama | m | -9.1525 | 78.689722 | SAMN05224190 |
| STRI_WOM_0088 | STRI_WOM_0088 | *Heliconius erato hydara* | Panama | m | -9.1525 | 78.689722 | SAMN05224197 |
| STRI_WOM_5193 | STRI_WOM_5193 | *Heliconius erato hydara* | Panama | m | -9.1525 | 78.689722 | SAMN05224200 |
| STRI_WOM_5351 | STRI_WOM_5351 | *Heliconius erato hydara* | Panama | m | -9.1525 | 78.689722 | SAMN05224201 |
| BC_0411 | BC0411 | *Heliconius erato lativitta* | Ecuador | m | -1.098333 | 77.583889 | SAMN05224101 |
| lativitta_01 | LAT01 | *Heliconius erato lativitta* | Ecuador | f | -1.098333 | 77.583889 | SAMN05224137 |
| lativitta_02 | LAT02 | *Heliconius erato lativitta* | Ecuador | f | -1.098333 | 77.583889 | SAMN05224138 |
| lativitta_03 | LAT03 | *Heliconius erato lativitta* | Ecuador | f | -1.098333 | 77.583889 | SAMN05224139 |
| lativitta_04 | LAT04 | *Heliconius erato lativitta* | Ecuador | f | -0.7125 | 77.583889 | SAMN05224140 |

| | | | | | | | |
|---|---|---|---|---|---|---|---|
| BC_0410 | BC_0410 | *Heliconius erato notabilis* | Ecuador | m | -1.81337 | -78.04507 | SAMN05224100 |
| BC_3223 | BC_3223 | *Heliconius erato notabilis* | Ecuador | m | -1.81337 | -78.04507 | SAMN05224105 |
| BC_3224 | BC_3224 | *Heliconius erato notabilis* | Ecuador | f | -1.81337 | -78.04507 | SAMN05224106 |
| BC_3225 | BC_3225 | *Heliconius erato notabilis* | Ecuador | f | -1.81337 | -78.04507 | SAMN05224107 |
| BC_3227 | BC_3227 | *Heliconius erato notabilis* | Ecuador | f | -1.82259 | -78.04406 | SAMN05224108 |
| BC_3228 | BC_3228 | *Heliconius erato notabilis* | Ecuador | m | -1.82259 | -78.04406 | SAMN05224109 |
| notabilis_01 | NOT01 | *Heliconius erato notabilis* | Ecuador | m | -1.399167 | 78.181111 | SAMN05224178 |
| notabilis_02 | NOT02 | *Heliconius erato notabilis* | Ecuador | m | -1.399167 | 78.181111 | SAMN05224179 |
| notabilis_03 | NOT03 | *Heliconius erato notabilis* | Ecuador | m | -1.399167 | 78.181111 | SAMN05224180 |
| notabilis_04 | NOT04 | *Heliconius erato notabilis* | Ecuador | m | -1.399167 | 78.181111 | SAMN05224181 |
| CA51 | CA51 | *Heliconius erato petiverana* | Mexico | f | 18.957903 | -90.269233 | SAMN05224115 |
| CA53 | CA53 | *Heliconius erato petiverana* | Mexico | m | 18.957903 | -90.269233 | SAMN05224116 |
| CA54 | CA54 | *Heliconius erato petiverana* | Mexico | m | 18.957903 | -90.269233 | SAMN05224117 |
| CA55 | CA55 | *Heliconius erato petiverana* | Mexico | f | 18.957903 | -90.269233 | SAMN05224118 |
| CA56 | CA56 | *Heliconius erato petiverana* | Mexico | m | 18.957903 | -90.269233 | SAMN05224119 |
| STRI_WOM_5732 | STRI_WOM_5732 | *Heliconius erato phyllis* | Bolivia | m | -18.176869 | -63.881664 | SAMN05224204 |
| STRI_WOM_5742 | STRI_WOM_5742 | *Heliconius erato phyllis* | Bolivia | m | -18.176869 | -63.881664 | SAMN05224205 |
| STRI_WOM_5765 | STRI_WOM_5765 | *Heliconius erato phyllis* | Bolivia | m | -18.176869 | -63.881664 | SAMN05224206 |
| STRI_WOM_5766 | STRI_WOM_5766 | *Heliconius erato phyllis* | Bolivia | m | -18.176869 | -63.881664 | SAMN05224207 |
| M_3654 | CS003654 | *Heliconius erato venus* | Colombia | m | 3.5311 | -76.753383 | SAMN05224141 |
| M_3655 | CS003655 | *Heliconius erato venus* | Colombia | f | 3.5311 | -76.753383 | SAMN05224142 |
| M_3656 | CS003656 | *Heliconius erato venus* | Colombia | m | 3.5311 | -76.753383 | SAMN05224143 |
| M_3657 | CS003657 | *Heliconius erato venus* | Colombia | m | 3.5311 | -76.753383 | SAMN05224144 |
| M_3659 | CS003659 | *Heliconius erato venus* | Colombia | m | 3.5311 | -76.753383 | SAMN05224145 |
| BC2565 | BC_2565 | *Heliconius himera* | Peru | f | -5.43724 | -78.4714 | SAMN08049963 |
| BC2566 | BC_2566 | *Heliconius himera* | Peru | m | -5.43724 | -78.4714 | SAMN08049964 |
| BC2567 | BC_2567 | *Heliconius himera* | Peru | m | -5.43724 | -78.4714 | SAMN08049965 |
| BC2570 | BC_2570 | *Heliconius himera* | Peru | f | -5.43724 | -78.4714 | SAMN08049966 |
| himera_001 | HIM001 | *Heliconius himera* | Ecuador | m | -4.276111 | -79.195833 | SAMN05224132 |
| himera_002 | HIM002 | *Heliconius himera* | Ecuador | m | -4.276111 | -79.195833 | SAMN05224133 |
| himera_003 | HIM003 | *Heliconius himera* | Ecuador | f | -4.276111 | -79.195833 | SAMN05224134 |
| himera_006 | HIM006 | *Heliconius himera* | Ecuador | f | -4.276111 | -79.195833 | SAMN05224135 |
| himera_030 | HIM030 | *Heliconius himera* | Ecuador | m | -4.276111 | -79.195833 | SAMN05224136 |
| Hermathena_13 | LMCI94-13 | *Heliconius hermathena* | Brazil | f | -2.453809 | -54.726839 | SAMN05224129 |
| Hermathena_14 | LMCI94-14 | *Heliconius hermathena* | Brazil | f | -2.453809 | -54.726839 | SAMN05224130 |
| Hermathena_15 | LMCI94-15 | *Heliconius hermathena* | Brazil | f | -2.453809 | -54.726839 | SAMN05224131 |

**Table S5. 2D *δaδi* model descriptions.**

| Model | Model details | Parameters |
|---|---|---|
| Split with complete isolation models | | |
| **SI** | Basic model with split and complete isolation. | *nu1, nu2, Ts, O* |
| **SI2N** | Model with split and complete isolation, heterogenous effective population size (2 classes, shared by the two populations = linked selection). | *nu1, nu2, Ts, nr, hrf, O* |
| **SIG** | Model with split, complete isolation and exponential growth. | *nu1, nu2, b1, b2, Ts, O* |
| **SI2NG** | Model with split, complete isolation, heterogenous effective population size (2 classes, shared by the two populations = linked selection) and exponential growth. | *nu1, nu2, b1, b2, hrf, Ts, Q, O* |
| *Divergence with migration models* | | |
| **IM** | Basic model with migration during divergence. | *nu1, nu2, m12, m21, Ts, O* |
| **IMG** | Model with migration during divergence and exponential growth. | *nu1, nu2, b1, b2, m12, m21, Ts, O* |
| **IM2N** | Model with migration during divergence and heterogenous effective population size (2 classes, shared by the two populations = linked selection). | *nu1, nu2, hrf, m12, m21, Ts, Q, O* |
| **IM2NG** | Model with migration during divergence, heterogenous effective population size (2 classes, shared by the two populations = linked selection) and exponential growth. | *nu1, nu2, b1, b2, hrf, m12, m21, Ts, Q, O* |
| **IM2m** | Model with migration during divergence and two types of migration. | *nu1, nu2, m12, m21, me12, me21, Ts, P, O* |
| **IM2mG** | Model with migration during divergence, two types of migration and exponential growth. | *nu1, nu2, b1, b2, m12, m21, me12, me21, Ts, P, O* |
| *Ancient migration models* | | |
| **AM** | Basic model with split and ancient migration. | *nu1, nu2, m12, m21, Ts, Tam, O* |
| **AMG** | Model with split, ancient migration and exponential growth. | *nu1, nu2, b1, b2, m12, m21, Ts, Tam, O* |
| **AM2N** | Model with split, ancient migration and heterogenous effective population size (2 classes, shared by the two populations = linked selection). | *nu1, nu2, hrf, m12, m21, Tam, Ts, Q, O* |
| **AM2N2m** | Model with split, ancient migration and heterogenous effective population size (2 classes, shared by the two populations = linked selection) and 2 ancient migration rates. | *nu1, nu2, hrf, m12, m21, me12, me21, Tam, P, Q, O* |
| **AM2NG** | Model with split, ancient migration and heterogenous effective population size (2 classes, shared by the two populations = linked selection) and exponential growth. | *nu1, nu2, b1, b2, hrf, m12, m21, Tam, Ts, Q, O* |
| **AM2m** | Model with split and 2 ancient migration rates. | *nu1, nu2, m12, m21, me12, me21, Ts, Tam, P, O* |
| **AM2mG** | Model with split, 2 ancient migration rates and exponential growth. | *nu1, nu2, b1, b2, m12, m21, me12, me21, Tam, Ts, P, O* |
| **AM2N2mG** | Model with split, ancient migration and heterogenous effective population size (2 classes, shared by the two populations = linked selection), 2 ancient migration rates and exponential growth. | *nu1, nu2, b1, b2, hrf, m12, m21, me12, me21, Tam, Ts, P, Q, O* |
| *Secondary contact models* | | |
| **SC** | Basic model with split, complete isolation, followed by secondary contact. | *nu1, nu2, m12, m21, Ts, Tsc, O* |
| **SC2N** | Model with split, complete isolation, followed by secondary contact and heterogenous effective population size (2 classes, shared by the two populations = linked selection). | *nu1, nu2, hrf, m12, m21, Ts, Tsc, Q, O* |
| **SCG** | Model with split, complete isolation, followed by secondary contact and exponential growth. | *nu1, nu2, b1, b2, m12, m21, Ts, Tsc, O* |
| **SC2N2m** | Model with split, complete isolation, followed by secondary contact, heterogenous effective population size (2 classes, shared by the two populations = linked selection) and 2 migration rates. | *nu1, nu2, hrf, m12, m21, me12, me21, Ts, Tsc, P, Q, O* |
| **SC2NG** | Model with split, complete isolation, followed by secondary contact, heterogenous effective population size (2 classes, shared by the two populations = linked selection) and exponential growth. | *nu1, nu2, b1, b2, hrf, m12, m21, Ts, Tsc, Q, O* |
| **SC2m** | Model with split, complete isolation, followed by secondary contact and 2 migration rates. | *nu1, nu2, m12, m21, me12, me21, Ts, Tsc, P, O* |
| **SC2mG** | Model with split, complete isolation, followed by secondary contact, 2 migration rates and exponential growth. | *nu1, nu2, b1, b2, m12, m21, me12, me21, Ts, Tsc, P, O* |
| **SC2N2mG** | Model with split, complete isolation, followed by secondary contact, heterogenous effective population size (2 classes, shared by the two populations = linked selection), 2 migration rates and exponential growth. | *nu1, nu2, b1, b2, hrf, m12, m21, me12, me21, Ts, Tsc, P, Q, O* |

**Table S 6. Bash pseudocode to run *msms* (Ewing and Hermisson 2010) simulations**. Note that mutation rate is only used by *seq-gen* when simulating the sequences, population size is used to scale times, selection and migration proportions.

```
for position in 0.5 -9.5 -19.5 -29.5 -39.5 -49.5 -59.5 -69.5 -79.5 -89.5 -99.5 -109.5 -119.5 -129.5 -139.5 -149.5 -159.5 -169.5 -179.5 -189.5 -199.5;
do for r in 0.000625 0.00125 0.0025 0.005 0.01 0.02 0.04 0.08;
do for m in 0.0000001 0.000001 0.00001;
do for t1 in 0.25 0.025 0.0025;
do for tm in 0.50000 0.25000 0.12500 0.02500 0.00250 0.00025 0.000025;
do for ts in 0.5 0.05 0.005 0.0005;
do

./msms 40 1 -N 1000000 -T -I 4 10 10 10 10 \
-Sp $position -r $r*10000 10000 \
-ej 0.025 1 2 -ej 0.5 2 3 -ej 1 3 4 \
-m 2 3 $m*4*1000000 -m 3 2 $m*4*1000000 -em $tm 2 3 0.0 -em $tm 3 2 0.0 \
-Smu 0.1 -SI $ts 4 0.0 0.0 0.01 0.01 \
-Sc 0 1 0.2*4*1000000 0.1*4*10000000 -Sc 0 2 0.2*4*1000000 0.1*4*10000000 0 -Sc 0 3 0 0.1*4*10000000 0.2*4*1000000 -Sc 0 4 0 0.1*4*10000000 0.2*4*1000000

done; done; done; done; done; done;
```

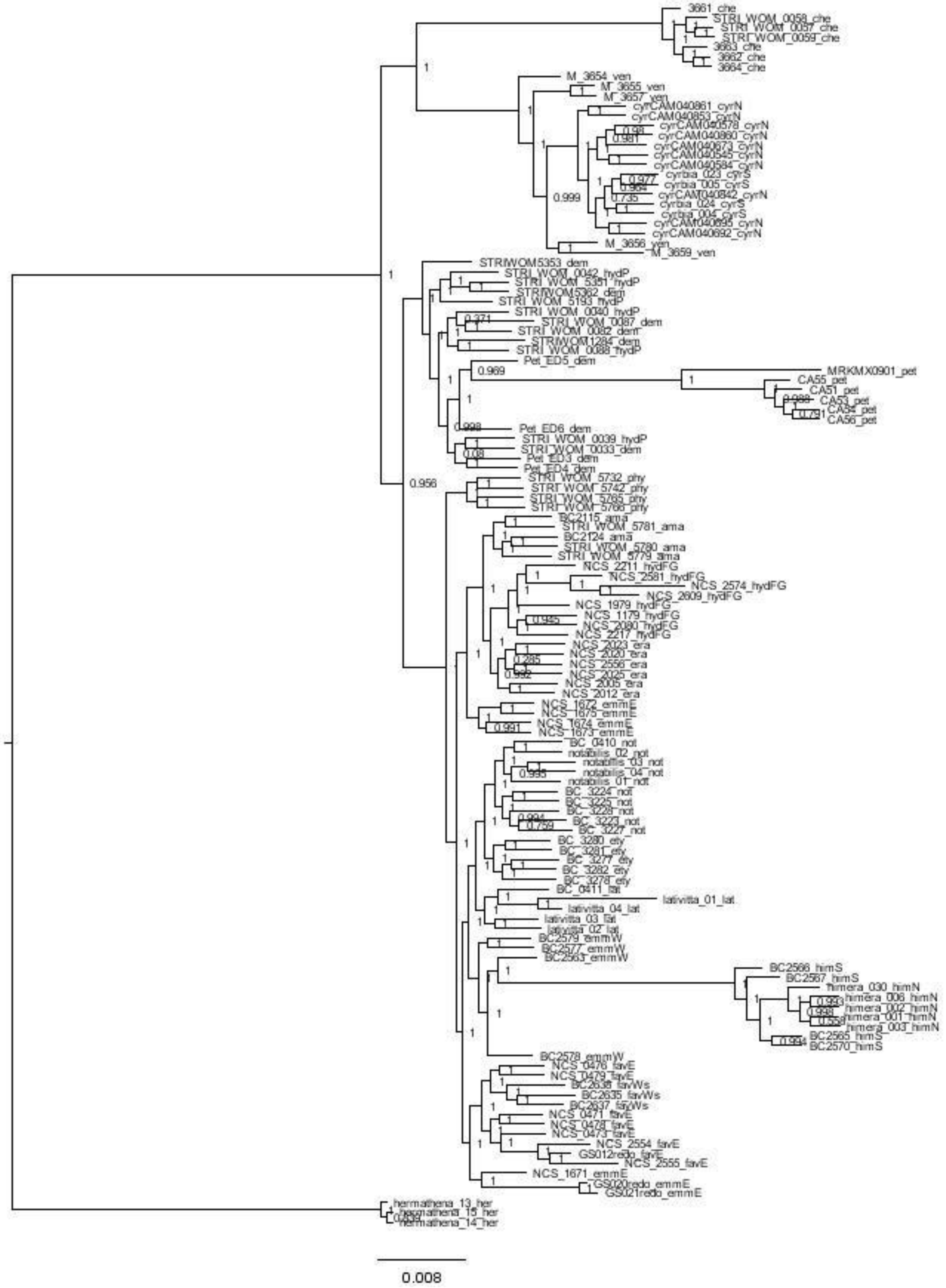

**Figure S 1. FastTree of samples used in this study.** Node support values are based on the Shimodaira–Hasegawa test.

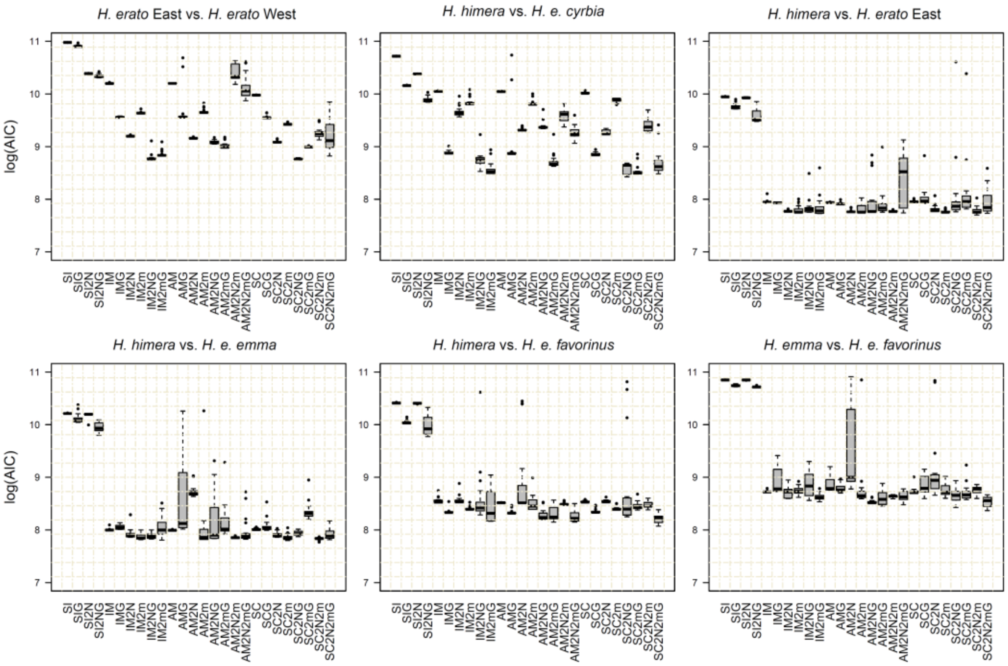


**Figure S 2. Average AIC (Akaike Information Criterion) of all tested *δaδi* models for twenty runs.** SI = Strict Isolation, IM = Isolation and Migration, AM = Ancestral Migration, SC = Secondary Contact, G = exponential growth, 2N = heterogenous effective population size (with 2 classes of loci shared by the two populations = Hill-Robertson effects), and 2m = 2 migration rates.

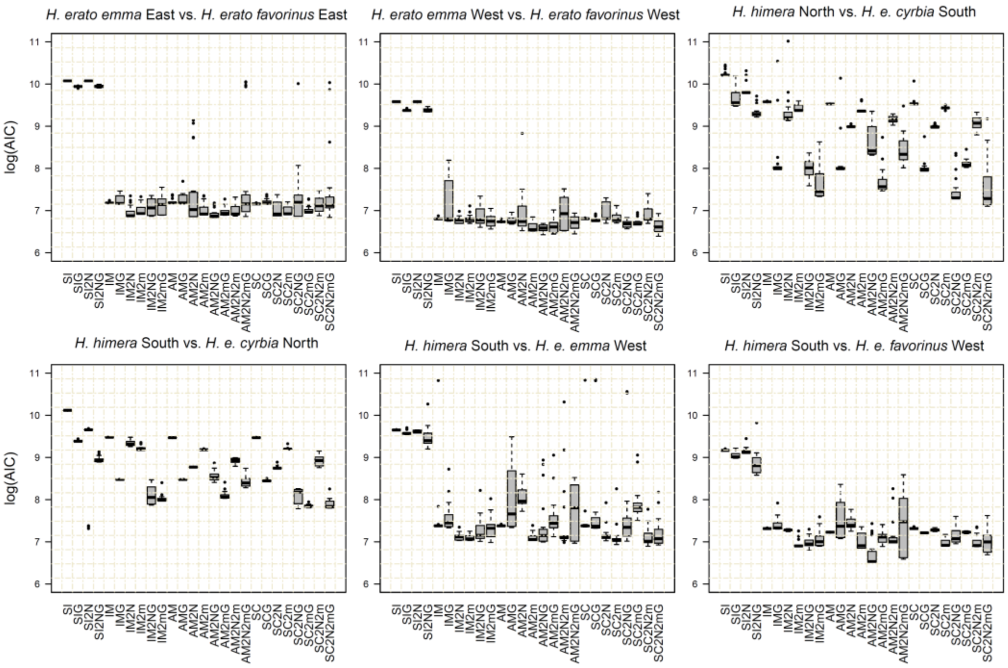


**Figure S 3. Average AIC (Akaike Information Criterion) of all tested *δaδi* models for twenty runs using subpopulations.** SI = Strict Isolation, IM = Isolation and Migration, AM = Ancestral Migration, SC = Secondary Contact, G = exponential growth, 2N = heterogenous effective population size (with 2 classes of loci shared by the two populations = Hill-Robertson effects), and 2m = 2 migration rates.

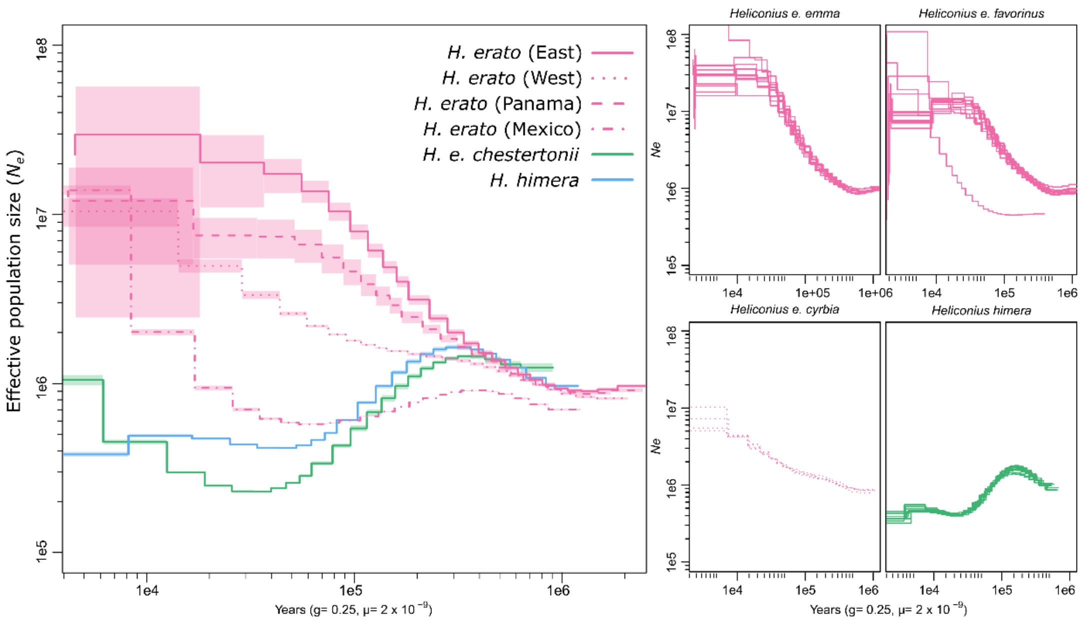


**Figure S 4. Inference of historical effective population size changes using pairwise sequentially Markovian coalescent (PSMC) analysis**. Lines and shading in the left panel represent the average effective population size and 95% CI, respectively, estimated from PSMC analyses on individual genome samples as shown in the panels on the right. The PSMC estimates are scaled using a generation time of 0.25 years and a mutation rate of 2e-9. Data adopted from Van Belleghem et al. (2018).

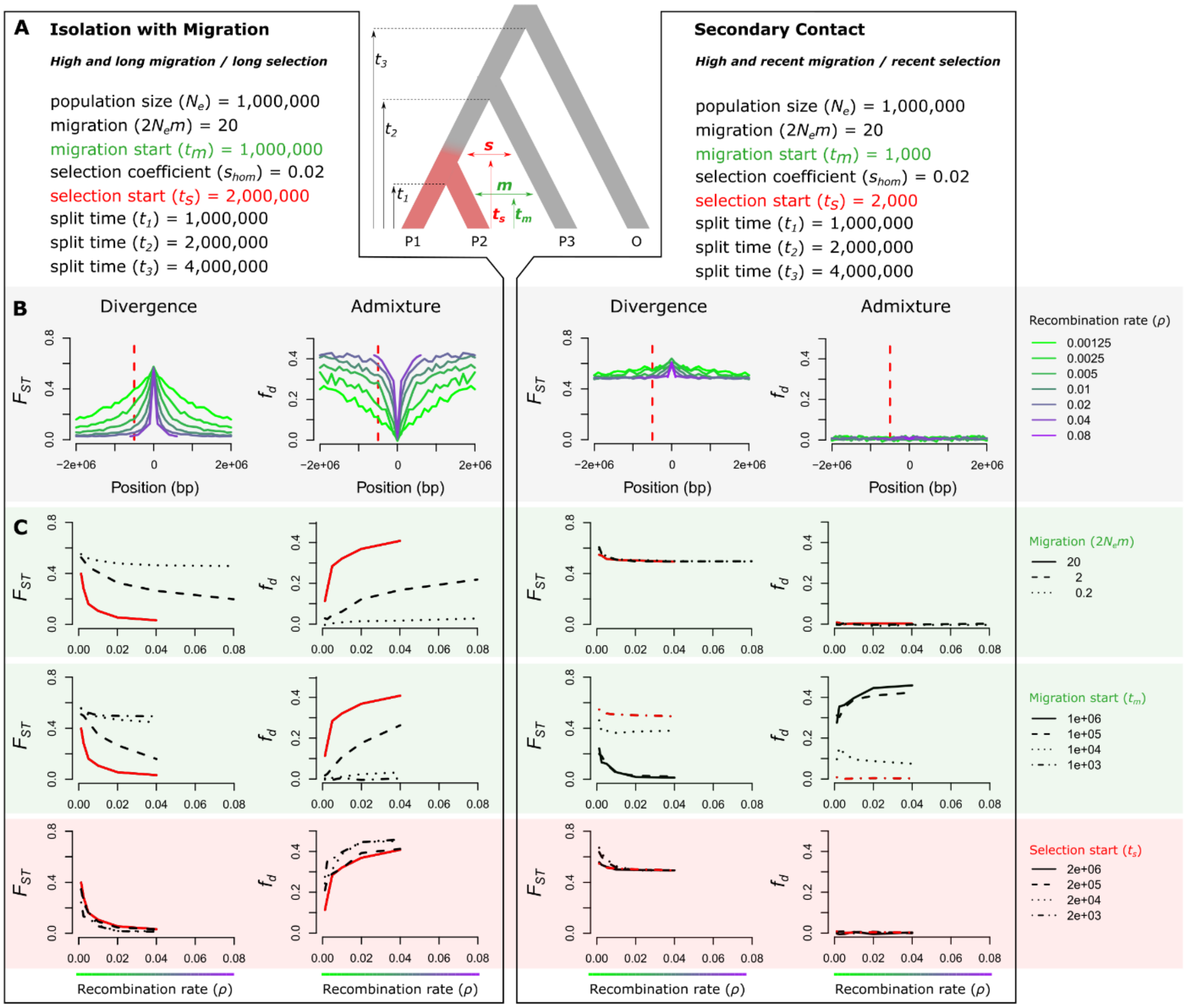


**Figure S 5. Expected relationship of recombination rate with divergence ($F_{ST}$) and admixture ($f_d$) near a divergently selected locus ($s$ = 0.02). (A)** The population tree shows the simulated scenario with the onset of divergent selection on a derived allele indicated in red and in which migration rate ($m$) and migration time ($t_m$) between P2 and P3 and selection start time ($t_s$) are varied. Left and right of the simulated scenario are parameter combinations for two extreme scenarios that both include linked selection; on the left a scenario with Isolation with Migration (IM) and on the right a scenario reflecting Secondary Contact (SC). **(B)** Effect of population recombination rate ($\rho$) on relative divergence ($F_{ST}$) and admixture ($f_d$) near a divergently selected locus for the two simulated scenarios with parameter combinations as in panel A. The selected allele occurs at position 0. The dashed red line indicates a locus at 500 kb from the selected locus at which the relationship between $\rho$, divergence and admixture is assessed in panel C. **(C)** The effect of migration start time ($t_m$) and selection start time ($t_s$) on the relation between $\rho$, divergence and admixture. Apart from the respective parameters being evaluated, other parameters were fixed as in panel A, with the red lines indicating the exact parameter combinations as in panel A and B.

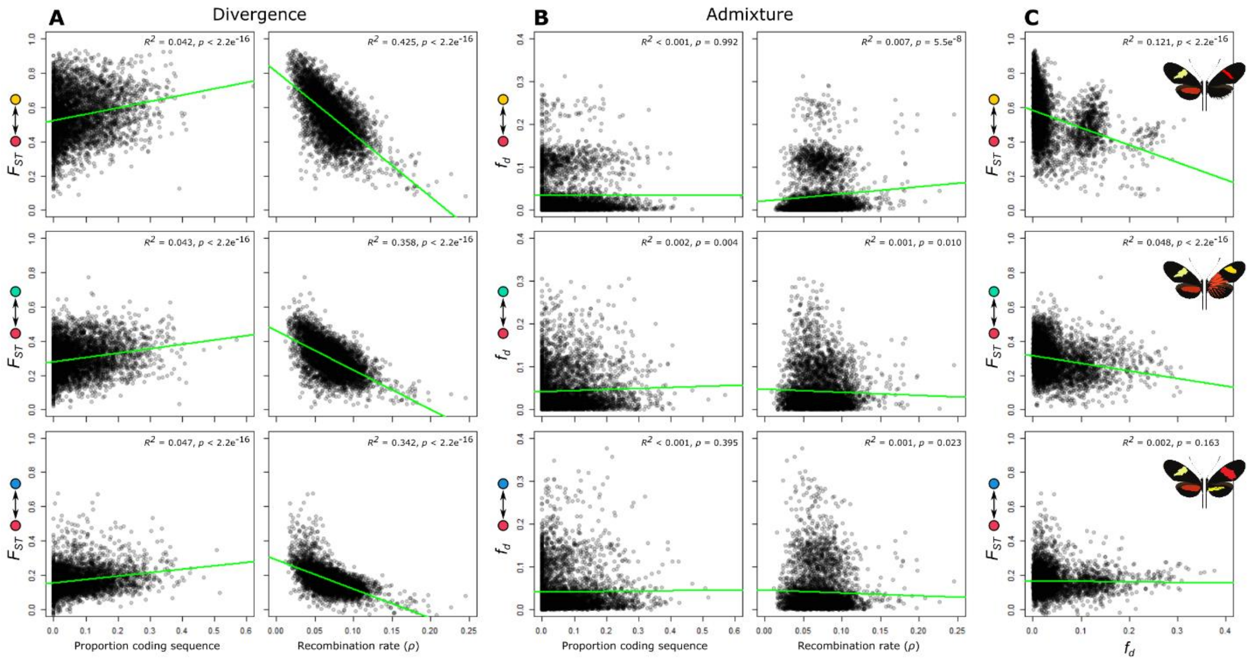


**Figure S 6. Correlations of divergence and admixture proportions with recombination rate and gene density in *H. himera* – *H. erato* contact zones. (A)** Relative divergence ($F_{ST}$) versus gene density and recombination rate ($\rho$). **(B)** Admixture proportion ($f_d$) versus gene density and recombination rate (cM/Mb). **(C)** Relative divergence ($F_{ST}$) versus admixture proportion ($f_d$). Statistics were calculated in 50 kb non-overlapping windows. Colored circles match color codes in Figure 2.
